# Structural Mechanisms Underlying Distinct Binding and Activities of 18:0 and 18:1 Lysophosphatidic acids at LPA1 Receptor

**DOI:** 10.64898/2025.12.09.693124

**Authors:** Ayobami Diyaolu, Peter Obi, Pravita Balijepalli, Kathryn Meier, Senthil Natesan

## Abstract

Lysophosphatidic acids (LPAs) are bioactive lipids that regulate numerous physiological functions in humans. Cell signaling by LPAs is mediated mainly via six LPA receptors (LPA1-6), class A G protein-coupled receptors (GPCRs). Among these, LPA1 is recognized to play an essential role in cell proliferation, survival, migration, and tumorigenesis. Despite the structural similarity, 18:0-LPA and 18:1-LPA exhibit distinct functional responses in cell lines overexpressing LPA1. Specifically, our in vitro studies show that 18:1-LPA induces greater Erk activation than 18:0-LPA in PC-3 human prostate cancer cells. The structural basis underlying this differential receptor activation has not been previously studied. Using classical molecular dynamics and enhanced sampling techniques, we examined the access and binding mechanisms of the two LPA species to the active state LPA1 receptor. The results show that 18:0-LPA and 18:1-LPA adopt distinct and dynamic poses in the orthosteric pocket despite their similar starting configurations. Mainly, the alkyl tails of the ligands exhibit distinct orientations and residue interactions, leading to differential conformational changes in key activation switches on the conserved CWxP and PIF structural motifs of the receptor. Also, there are significant differences in interhelical interactions at the intracellular end of the transmembrane helices 1, 3, 6, and 7. These distinct arrangements lead to striking differences in LPA1 interactions with the Gα-helix of the heterotrimeric Gi-protein. Notably, 18:0-LPA and 18:1-LPA exhibit similar membrane partitioning characteristics and receptor entry processes through aqueous paths. Our comprehensive in-silico studies offer valuable structural insights into the observed differences in functional responses by 18:0-and 18:1-LPA.

## INTRODUCTION

Lysophosphatidic acid (LPA) receptors belong to a subfamily of class A G protein-coupled receptors (GPCRs) known as Lipid GPCRs. Lipid GPCRs also include sphingosine 1-phosphate receptors and cannabinoid receptors and are named based on the lipid-derived origins of their endogenous agonists^1,2^. The LPA receptor family includes six receptors (LPA 1-6), among which LPA1 is the most studied^3, 4^. LPA1 is widely distributed in human tissues, and its activation promotes cell proliferation, survival, migration, and tumorigenesis^5–7^. Notably, LPA1 has been implicated in cancer cell survival and nourishment of the tumor microenvironment^6^. LPA1 is promiscuous regarding G-protein coupling^8^. It can couple to the G_12/13_ protein to activate phospholipase C, which leads to calcium mobilization^5, 9^. It can also bind to Gi/o protein, through which it modulates the RAS pathway, ultimately leading to the phosphorylation of the extracellular signal-regulated kinase (Erk)^10^.

LPA1 is activated by endogenous lysophospholipids known as lysophosphatidic acids (LPAs). LPAs are nearly ubiquitous in human body tissues and fluids^7, 11^. Structurally, they possess a phosphate head, glycerol backbone, and an alkyl tail that varies in length and degree of unsaturation, giving rise to numerous LPA species^12, 13^. The most abundant LPA species in the human body include 14:0, 17:0, 18:0, 18:1, and 20:4 LPAs^14^. The relative activities of various forms of LPA have been reported for a handful of cellular models in which the relevant LPA receptor was defined. In 2000, Bandoh and colleagues examined the activities of an extensive series of LPA analogs on calcium mobilization in Sf9 insect cells transfected with LPA1, LPA2, or LPA3 receptors^15^. They reported similar potencies for 18:1-, 18:2-, 18:3-, and 20:4-LPAs in cells expressing LPA1 receptor. There was negligible agonist activity for 12:0- and 14:0-LPAs and lower potency for 16:0- and 18:0-LPAs than for 18:1-LPA. Subsequently, another study examined the effects of multiple LPA species on calcium mobilization and Erk activation in human lung fibroblasts, where LPA1 is the predominant endogenous LPA receptor^10^. With respect to Erk activation, the study reported that 20:0-, 20:4-, 18:2-, 18:3-, 17:0- 16:0-, and 14:0-LPAs had lower potency than 18:1-LPA. Also, longer chain LPAs (18:1 and 20:4) have shown greater activity than shorter chain LPAs (14:0 and 16:0)^10, 14^. This study also reported possible bias between calcium mobilization and Erk activation for some ligands (16:0, 17:0, and 18:2), suggesting that relative efficacy can depend on the measured response.

We previously showed that 18:1-LPA was most effective at increasing Erk activation in Du145 prostate cancer cells, 18:0-LPA acted as a partial agonist, and 16:0-, 14:0-, and 6:0-LPAs elicited little or no response^16^. Subsequent work by our group confirmed that LPA1 is the receptor responsible for 18:1-LPA-induced proliferation and migration in Du145 and PC-3 human prostate cancer cells^17^. In summary, previous findings indicate that 18:1-LPA is the most potent and efficacious agonist for the LPA1 receptor. Shorter saturated LPA species and other 18-carbon LPAs with additional saturation/unsaturation have varying efficacies in different model systems but are generally less active than 18:1-LPA. Data interpretation can be complicated by the fact that most mammalian cells express multiple LPA receptors since the relative activities of different LPA species differ between LPA receptors^3, 15, 18^.

18:1-LPA is nearly identical to 18:0-LPA in structure and almost every physicochemical property; the only difference is the presence of a double bond between C9 and C10 carbons in 18:1-LPA, whereas 18:0-LPA is fully saturated **(Figure 1A and Table S1)**^19^. Both molecules are amphiphilic, with 18:0-LPA (Clog *P* = 5.85) slightly more lipophilic than 18:1-LPA (Clog *P* = 5.49)^20^. The two LPAs are highly flexible due to their many rotatable bonds (18:1-LPA = 21, 18:0-LPA= 22) and thus can potentially exist in many conformations under physiological conditions. Despite their similarities,18:1-LPA is a more efficacious agonist for LPA1 than 18:0-LPA, as introduced earlier. The difference in the saturation of the alkyl chain may give rise to the differential behavior of the two species in lipid bilayers and in their interactions with the membrane-embedded receptors. In general, saturation in fatty alkyl chains of phospholipids has been shown to impact the biophysical characteristics of the membrane, including fluidity, bilayer thickness, curvature, and stability, which could eventually affect important cellular and biological functions^21, 22^. Although the role of fatty alkyl chain length on the activity of LPAs on lipid GPCRs has been studied before^23^, to our knowledge, no studies have examined the mechanistic basis for the impact of saturation-unsaturation on the activities of LPA species at LPA1.

**Figure 1.**
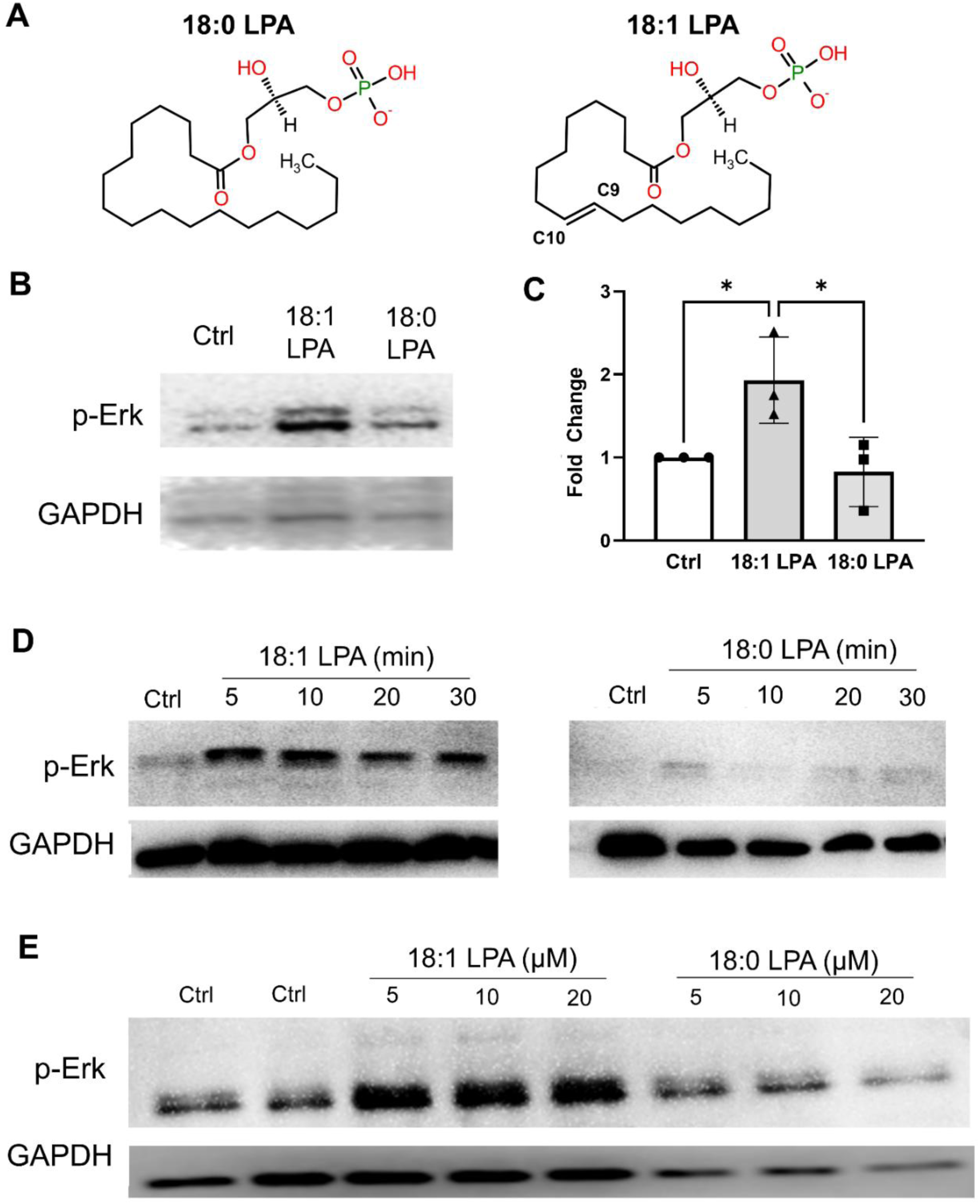
Effects of 18:1- and 18:0-LPA on Erk activation in PC-3 human prostate cancer cells. **A)** 2D structures of 18:0-LPA and 18:1-LPA. Both molecules are structurally identical, with a polar glycerol-phosphate group at one end and an 18-carbon alkyl chain at the other. The only difference is the double bond between C9 and C10 atoms in 18:1-LPA, which is absent in 18:0-LPA. **B)** Serum-starved PC-3 cells were incubated with and without (“Ctrl”) 10 µM 18:1-LPA and 18:0-LPA for 15 minutes.

Recent developments in structural biology (cryo-EM and X-ray crystallography) led to the elucidation of the inactive LPA1 receptor bound to antagonists in 2015^24^. However, it wasn’t until recently that the active structure of LPA1 bound to endogenous 18:1-LPA was elucidated^25^. Subsequent developments in the field led to several active LPA1 receptor structures with co-crystallized lipid and non-lipid agonists^26, 27^. Compared to the inactive receptor, several structural rearrangements occur in the orthosteric pocket of active LPA1 involving several hydrophobic residues, including L132^3.36^, W210^5.43^, F267^6.44^, W271^6.48^, L297^7^.39, and A300^7.42 24^. LPA1 receptor activation involves a large outward movement of the intracellular end of transmembrane helix 6 (TM6) and an inward movement of TM7 toward TM3^28^. These changes lead to the opening of LPA1’s intracellular cavity for coupling to the Gα-helix of the Gi protein. Similar to other Class A GPCRs^28–31^, signal transduction between the binding site and G-protein coupling interface could involve conserved structural domains, including the CWxP motif (C^6.47^, W^6.48^, x^6.49^, and P^6.50^), PIF motif (P^5.50^, I^3.36^, and F^6.44^), and NPxxY motif (N^7.49^, P^7.50^, x^7.51^, x^7.52^, and Y^7.53^). However, most of these structural details on LPA1 activation by agonists have been based majorly on the static structures from cryo-EM studies. There is limited information on comparative changes in the conformational dynamics of active state LPA1 in the presence of endogenous agonists, which can be investigated via molecular dynamics (MD) simulations.

In addition, the mechanistic details of how 18:0-LPA and 18:1-LPA access the orthosteric binding site of LPA1 remain poorly understood. Although many of the endogenous ligands enter their orthosteric sites in GPCRs from the extracellular aqueous phase, for multiple receptors, ligand entry through transmembrane helices from the surrounding membrane lipids has been reported^32–34^. The mode of entry of LPA species could play a role in “pre-organizing” ligands into orientations and conformations that would affect the binding kinetics and interactions within the binding site^12, 35^. The polar head groups and hydrophobic alkyl tails of the LPA species should exhibit distinct affinities for the aqueous phase and various polar and nonpolar functional groups of the membrane lipids. These differences could affect their partitioning characteristics, such as preferred bilayer depth, orientation, and conformations. The membrane partitioning characteristics of these ligands are yet to be examined and correlated with their access and binding mechanisms to the orthosteric site of LPA1.

In this study, we first investigated the efficacy of 18:0-LPA and 18:1-LPA in activating Erk in PC-3 human prostate cancer cells. LPA1 is the endogenous receptor predominantly responsible for LPA-induced proliferation in this cell line^36^. Next, to gain structural insights into the activation process, we performed unbiased atomistic simulations of both 18:0 and 18:1-LPA bound to LPA1 in the presence and absence of the heterotrimeric Gi protein. Subsequently, we elucidated the plausible access paths of the ligands to the orthosteric site of the LPA1 receptor using well-tempered metadynamics (WT-metaD). Also, we utilized steered MD and umbrella sampling techniques to elucidate the membrane partitioning characteristics of both LPA species to determine their energetically favorable bilayer locations, orientations, and conformations within a model membrane bilayer. In addition, we computed the relative binding free energies of 18:0-LPA and 18:1-LPA to LPA1, including the contribution of individual binding site residues using the molecular mechanics Poisson-Boltzmann surface area (MM-PBSA) technique. Comprehensive analyses of both classical and enhanced sampling simulations revealed distinct and dynamic binding orientations and critical residue interactions of the two LPA species within the orthosteric site. The similarities and differences in hydrophobic interactions, rotameric shifts, several activation signatures commonly observed in other GPCRs, and residue-residue communication between the binding site residues and G protein coupling interface residues were analyzed to gain valuable insights into the differential activities of the two LPA species.

## RESULTS

### 18:1-LPA is more efficacious than 18:0-LPA at activating Erk in PC-3 human prostate cells

PC-3 human prostate cells respond to 18:1-LPA by increasing proliferation and migration; these responses are mediated by the LPA1 receptor^36^. Erk activation, which is involved in mitogenesis, is an early response to 18:1-LPA in PC-3 cells ^16, 37^. We, therefore, used LPA-induced Erk activation to compare the relative activities of 18:1- and 18:0-LPA in these cells. As shown in **Figure 1B**, 18:1-LPA activated Erk (as assessed by an antibody recognizing phosphorylated Erk) to a greater extent than 18:0-LPA when both ligands were applied at a concentration of 10 µM for 15 minutes. This response was quantified by densitometry from multiple experiments (**Figure 1C**). The results show that Erk phosphorylation in response to 18:1-LPA was statistically different from untreated cells and 18:0-LPA-treated cells; 18:0-LPA did not significantly increase Erk phosphorylation. These differences in efficacy between 18:1- and 18:0-LPA are consistent with those published previously for Du145, another human prostate cancer cell line ^16^, except that 18:0 was a partial agonist in Du145 cells. The relative efficacy is also consistent with the induction of cellular communication network (CNN) proteins, an alternative and more delayed LPA response observed in our study^38^. A time course experiment was carried out to test whether the different efficacies of 18:1- and 18:0-LPA reflected differences in the kinetics of the response (**Figure 1D**). The results show that 18:0-LPA did not elicit substantial Erk activation at times from 5-30 minutes, as compared with the response to 18:1-LPA, which was maximal at 10 minutes but prominent at all time points tested. Next, the effects of different concentrations of 18:1- and 18:0-LPA were compared to determine whether 18:0 might be less potent than 18:1-LPA (**Figure 1E**). The results indicate that the response to 18:0-LPA was negligible at all concentrations tested (5-20µM), while 18:1-LPA elicited Erk activation throughout this dose range. In summary, the signal transduction studies, which were conducted at early times after LPA addition, indicate that 18:1-LPA is an agonist for PC-3 cells expressing endogenous LPA1, while 18:0-LPA has negligible efficacy.

Whole-cell extracts were immunoblotted for activated phospho-Erk (p-Erk) and for GAPDH (loading control). **C)** Erk phosphorylation was quantified from experiments in which serum-starved PC-3 cells were incubated with and without 10 µM 18:1-LPA and 18:0-LPA for 10 minutes. The p-Erk signal was quantified by densitometry, utilizing background subtraction and normalization to GAPDH. The p-Erk levels in treated cells were divided by levels in untreated control cells within each experiment. Each data point represents the mean ± S.D. of values from four separate experiments. Statistical significance was determined using the Student’s t-test. **D)** Serum-starved PC-3 cells were incubated with 10 µM 18:1-LPA or 18:0-LPA for 5-30 minutes. Whole-cell extracts were immunoblotted for p-Erk and GAPDH. All lanes are from the same experiment, gel, and blot, except that p-Erk and GAPDH were analyzed from separate gels using the same samples. **E)** Serum-starved cells were incubated for 10 minutes with the indicated concentrations of 18:1- or 18:0-LPA; the control (“Ctrl”) was untreated. Whole-cell extracts were immunoblotted for p-Erk and GAPDH.

### Dynamic binding modes of LPAs in the LPA1’s orthosteric site

To assess the differential ligand binding and conformational dynamics of LPA1 upon binding to the two LPAs, we utilized a recently published cryo-EM structure of LPA1-Gi protein complex bound to 18:1-LPA (PDB ID: 7TD1). The receptor is illustrated in its secondary structure representation with critical orthosteric binding site residues and residues involved in signal transduction and activation processes (**Figure 2A**). In the static structure, the alkyl tail end of 18:1-LPA adopts a nearly U-shaped orientation within the pouch-shaped pocket created by steric restraints from W210^5.43^ at the base of the binding site^25^ **(Figure S1A)**. The ligand’s alkyl chain is also in contact with residues D129^3.33^, L132^3.36^, W271^6.48^, and L297^7.39^, lining the pocket and is oriented towards the space between TM1 and TM7. Since no experimental structure of LPA1 with 18:0-LPA is available, we performed several rounds of molecular docking to obtain the plausible binding poses. Ligand poses with the best docking scores (ranging from -12 to -15 kcal/mol) had the phosphate headgroup oriented towards the N-terminal interacting with residues Y34^Nterm^ and K39^Nterm^, similar to the 18:1-LPA bound structure. At the same time, the alkyl chain forms numerous hydrophobic interactions with residues lining the base of the orthosteric pocket **(Figure S1B)**. Overall, the selected pose of 18:0-LPA is within 4 Å of several binding site residues, including T109^ECL1^, T113^ECL1^, R124^3.28^, Q125^3.29^, D129^3.33^, W210^5.43^, W271^6.48^, E293^7.35^, and K294^7.36^, as previously described for 18:1-LPA^26^.

**Figure 2.**
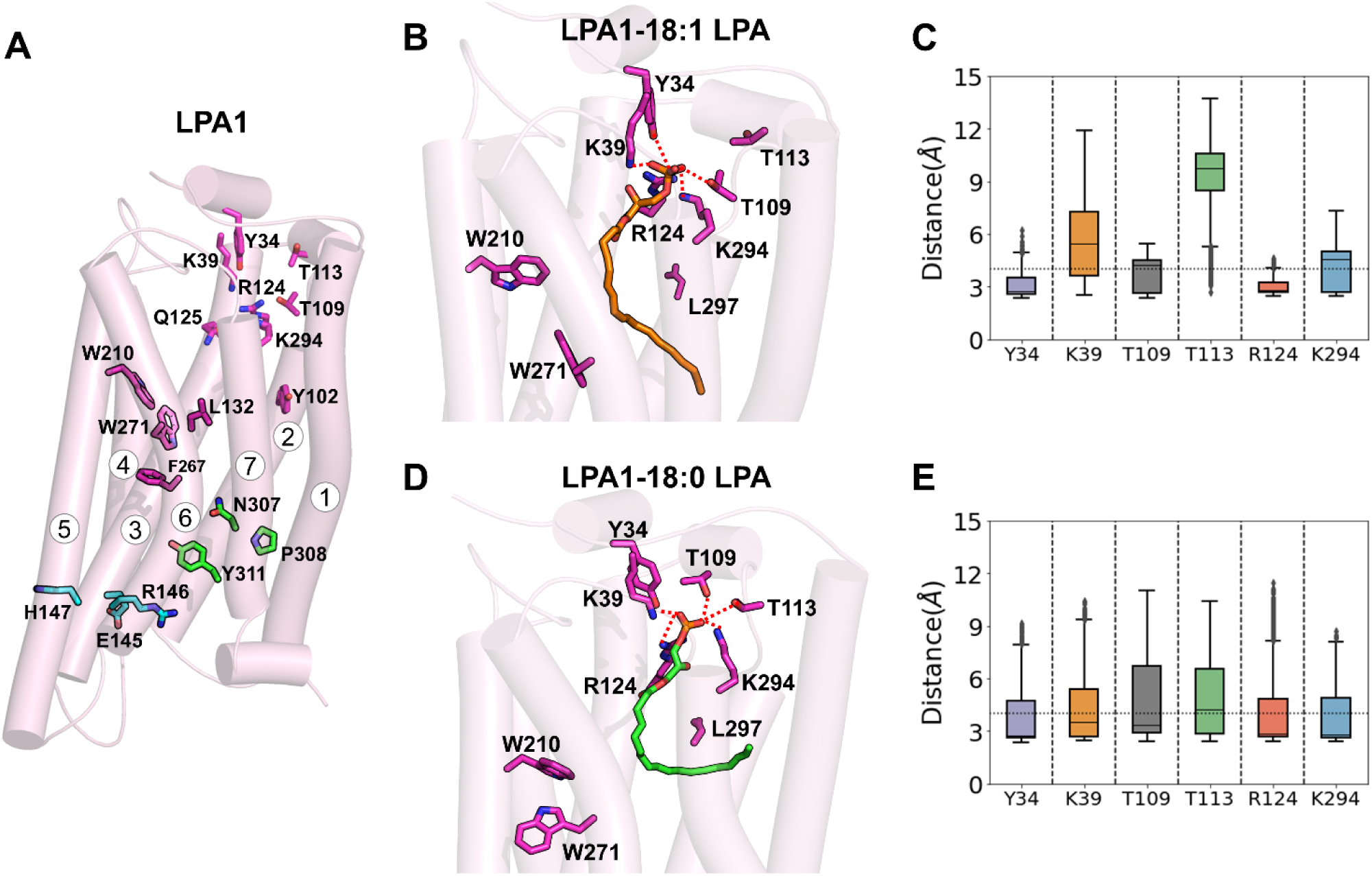
Binding poses and critical residue interactions of the LPA species observed in unbiased MD simulations. (A) The secondary structure representation of the LPA1 receptor in its active state (PDB ID 7TD1) with its seven transmembrane helices labeled 1-7. The sidechain groups of the orthosteric binding site residues and those involved in the access and binding of LPA ligands at the upper end of the receptor, residues involved in binding and signal transduction, and those involved in activation processes are highlighted in light magenta, dark magenta, green, and cyan colors, respectively. (B) The most preferred binding orientation of 18:1-LPA (orange licorice) within the orthosteric binding pocket was obtained from 500 ns MD simulations. The polar interactions of the ligand with several critical ECL1 and site residues are shown in dotted red. (C) The dynamics and stability of several polar interactions (salt bridges, H-bond, and other electrostatic interactions) by the glycerol-phosphate groups of 18:1-LPA with the binding site residues Y34^Nterm^, K39^Nterm^, T109^ECL1^, T113^ECL1^, R124^3.28^, and K294^7.36^ are shown as a distance box plot with their medians and quartiles. Similarly, the preferred binding pose, critical interactions, and distances for 18:0-LPA (green licorice) are shown in (D) and (E), respectively.

To investigate the dynamics of the ligands within the receptor site, we performed classical MD simulations for 500 ns (three replicates) of the LPA1-ligand receptor complexes in the presence and absence of the heterotrimeric Gi-protein. Altogether, three microsecond-long simulations were performed for each ligand (18:0- and 18:1-LPA). We observed consistent interactions and evolution of ligand poses over time in all three replicates. Therefore, results from representative simulations are used to describe the ligand binding dynamics here **(Figure 2)**. The longevity of polar and nonpolar residue-ligand interactions was quantified as contact frequency and presented as a heatmap (**Figure 3A**). The contact frequency represents the fraction of the simulation time during which a given binding site residue is within 4 Å of the ligand atoms.

**Figure 3.**
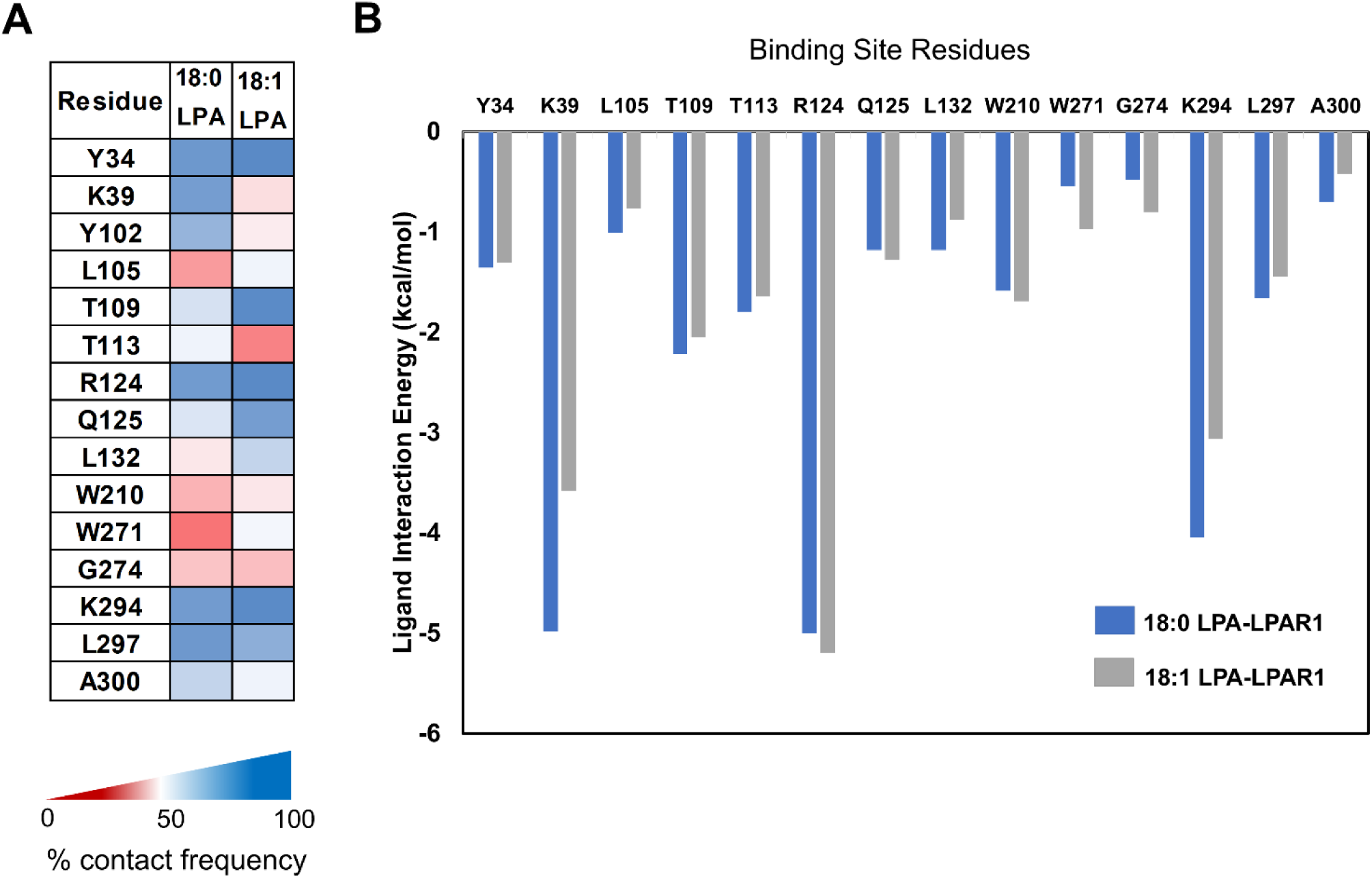
Binding site residue contacts and interaction energies of the LPA ligands with the active state LPA1 receptor. **(A)** The heatmap depicts the binding site residues’ contact frequency with 18:0-LPA and 18:1-LPA. The contact frequency represents the percentage of the simulation time during which a given residue is within 4 Å of the ligand. **(B)** The bar graph shows the relative contribution of specific LPA1 binding site residues to the estimated binding energies of 18:0-LPA (blue) and 18:1-LPA (grey).

The MD simulations primarily reveal significant conservation of polar contacts for both LPA species. The negatively charged phosphate group of 18:1-LPA engages in a salt-bridge interaction with the positively charged sidechain amino group of K39^Nterm^ residue at the N-terminal end of the receptor. This charge-charge interaction remains stable for the first 250 ns (**Figures 2B, 2C, and S1C**). The phosphate group also engages in polar H-bond interaction with the sidechain -OH group of Y34, which remained stable throughout the simulation time (500 ns). In contrast, in the case of 18:0-LPA, although there was a transient breakage of the salt bridge interaction with K39 around ∼100-250 ns, the interactions remained intact for the remainder of the simulation **(Figure 2D and S1D)**. Additionally, there were H-bond interactions between LPAs with extracellular loop (ECL1) residues T109^ECL1^ and T113^ECL1^ of LPA1. The interaction with T109^ECL1^ was stable for 18:1-LPA throughout the simulation, with a high occupancy rate. However, slight loop movements led to a change in the conformation of T113^ECL1^, resulting in breakage of contact from 18:1-LPA at approximately the first 10 ns **(Figure 2C & S1C)**. In contrast, 18:0-LPA had much more stable H-bonds with both T109^ECL1^ and T113^ECL1^ persisting for ∼350 ns, eventually re-orienting away from the ECL1 loop towards the end of the simulation **(Figure 2D & 2E)**.

The glycerol-phosphate groups of the two LPAs also made critical interactions within the transmembrane helices. The phosphate group oxygen atoms in 18:0- and 18:1-LPA were involved in salt bridge interactions with R124^3.28^ and K294^7.36^ from TM3 and TM7, respectively, with above-average occupancy rates **(Figures 2C & 2E)**. This is unsurprising as mutation of both these residues has been shown to disrupt ligand binding^39^.

Due to the flexibility of the two LPA species, they sampled multiple conformations away from the starting pose during the simulation. We measured the ligand internal angle between the phosphate (P1) head, the middle of the alkyl chain (C9-C10), and the tip of the alkyl chain (C1) **(Figure 1A & S1E)**. For 18:1-LPA, an initial internal angle of ∼60° and a U-shaped nature of the alkyl chain evolved to a more extended conformation with an internal angle of ∼105° toward the end of the simulation **(Figure S1E)**. This extended conformation gave the alkyl chain a larger surface for hydrophobic interactions with the pocket lining residues. While 18:0-LPA also sampled some conformational stretching, as marked by a transient increase in the internal angle, its alkyl chain eventually adopted a nearly L-shaped conformation (internal angle of ∼75°) toward the end of the simulation. Near the end of the simulation (∼400 ns), the lower end of the 18:0-LPA alkyl chain points outwardly through the space between the TM1 and TM7 towards the membrane bulk **(Figure 2C)**. As a result, the ligand moved toward TM3 and TM7, reducing its interactions with the ECL1 **(Figure S1D)**. In contrast, 18:1-LPA, with conformational restraints due to its double bond, can squeeze between the residues lining the base of the pocket without extending toward the membrane. The analysis of ligands’ pose stability showed RMSD values, on average, above 5 Å for both 18:0- and 18:1-LPA due to the high flexibility of the alkyl chain **(Figure S1F)**. The extra displacement of 18:0-LPA towards TM7 could partially account for its higher RMSD (7.7±2.3 Å) compared to 18:1 LPA (5.7±1.1 Å) **(Figure S1F)**. Overall, 18:1-LPA was more stable at the LPA1 orthosteric site than 18:0-LPA. Similar residue-ligand interaction patterns were observed in simulations without Gi protein (**Figure S4**).

### Distinct hydrophobic interactions at the base of the binding pocket

In addition to the strong charge-charge and polar-charge interactions, the ligands exhibit many polar and non-polar contacts within the binding pocket. To understand how these contacts evolve during the entire simulation, we computed contact frequency of residues within 4 Å of various parts of each ligand, including the glycerol-phosphate head and the alkyl chain. Overall, both ligands interact extensively with the N-terminus, TM2, ECL1, TM3, TM6, and TM7, although each ligand has its unique contact fingerprint at each region. Within the N-terminus, 18:0- and 18:1-LPA exhibit a high contact frequency with Y34^Nterm^. However, 18:1-LPA has much less contact with K39^Nterm^ than 18:0-LPA. There was a similar trend in TM2, where 18:0-LPA has greater contact with Y102^2.57^ than 18:1-LPA, which has better contact with L105^2.60^ **(Figure 3A)**. In addition, within the ECL1 region, 18:0-LPA is in more frequent contact with T113^ECL1^, while 18:1-LPA prefers interaction with T109^ECL1^. Interestingly, both species have similar overall contact profiles with TM7 residues, including G274^6.51^, K294^7.36^, L297^7.39^, and A300^7.42^ **(Figure 3A)**.

Except for the differential contact discussed earlier for the N-terminus and TM2, the glycerophosphate groups of both the ligands have similar contact profiles with residues in TM3 (W121^3.25^, R124^3.28^, Q125^3.29^), ECL2 (M198^ECL2^, P200^ECL2^) and TM7 (E293^7.35^ and K294^7.36^). This is unsurprising, as both ligands’ head groups are identical **(Figure S2A)**. However, the alkyl chain contacts in both ligands show remarkable differences. The 18:0-LPA alkyl chain drifted more towards the TM1-TM2 during the simulation, leading to contacts with V59^1.42^, C60^1.43^, and I63^1.46^. These contacts are reduced or absent in the case of 18:1-LPA **(Figure S2A)**. Consequently, 18:0-LPA has a higher contact with TM2, as noted previously for the A98^2.53^ and Y102^2.57^ interactions. In contrast, 18:1-LPA shows greater contact with TM3 residues, including Q125^3.29^, D129^3.33^, and L132^3.36^, which have all been shown to be important for agonist activity. The 18:1-LPA alkyl chain also has slightly higher contact with W210^5.43^, which forms a part of the pocket base, limiting the extension of the ligand deep down into the receptor **(Figure S2)**. At the TM6 region, the alkyl chains of both ligands have similar contacts with G274^6.51^.

However, 18:1-LPA preferentially contacts W271^6.48^ of the C^6.47^W^6.48^xP^6.50^ motif, unlike 18:0-LPA, which interacts with P273^6.50^ of the same motif **(Figure S2)**. Both 18:0 and 18:1-LPA alkyl chains make extensive hydrophobic contacts with residues in TM7, especially F296^7.38^, L297^7.39^, and A300^7.42^, with only subtle differences as outlined in the contact fingerprint plot **(Figure S3A-B)**.

Subsequently, we utilized the MM-PBSA method to evaluate the relative contribution of individual orthosteric site residues to the binding free energy of the two ligands. Generally, residues having polar interactions with the ligands, including K39^Nterm^, T109^ECL1^, T113^ECL1^, R124^3.28^, and K294^7.36^, contributed more significantly to the relative binding energy of both ligands compared to hydrophobic residues **(Figure 3B)**. These observations are due to the stable and consistent nature of the polar and charged interactions, as discussed earlier, based on the ligand’s hydrogen bond distances and contact frequencies. However, the hydrophobic residues (L105^2.60^, L132^3.36^, W210^5.43^, and W271^6.48^) exhibited more significant on-off contact dynamics, hence their lower contribution to ligand affinity. Particularly, R124^3.28^ and K294^7.36^ showed the highest contribution to both 18:0 and 18:1 LPA binding affinity. The importance of this residue has also been evaluated through mutagenesis studies^25^. K39^Nterm^ has a higher binding affinity for 18:0-LPA compared to 18:1-LPA, which correlates with the higher contact of the residue with 18:0 LPA **(Figure 3B)**. Among the hydrophobic residues, L297^7.39^ had the highest contribution to 18:0 LPA (-1.62 kcal/mol) binding energy, while W210^5.43^ contributed more to 18:1 LPA (-1.67 kcal/mol) binding. Overall, both ligands exhibited high relative binding energy with slightly higher affinity for 18:0 LPA (-65.6 ± 4.2 kcal/mol) compared to 18:1 LPA (-63.9 ± 4.6 kcal/mol) **(Table S2)**. Hence, the greater agonist activity seen in 18:1 LPA is unlikely related to differences in ligand binding affinity.

### Insights on differential activation from dynamics of conserved motifs

Class A GPCRs have several conserved structural motifs and activation switches that exhibit marked differences upon activation due to ligand binding^28, 40, 41^. These motifs play critical roles in the transduction of ligand-initiated responses through a coordinated pattern of conformational changes and breakage or formation of specific interactions. Like other class A GPCRs, previous structural elucidation of both active and inactive structures of LPA1 has shown an outward movement of TM6 upon activation, while TM7 shifts inwardly towards TM3 **(Figure 4A)**. As a result, Y311^7.53^ from the N^7.49^P^7.50^xxY^7.53^ conserved motif comes in close contact with I142^3.46^ at the intracellular ends of TM3, moving away from V260^6.37^ located on TM6. We assessed the movement and interactions of the three helices (TM3, TM6, and TM7) by quantifying the distances between the three residues during the entire simulation time in all replicates **(Figure 4B-E)**.

**Figure 4.**
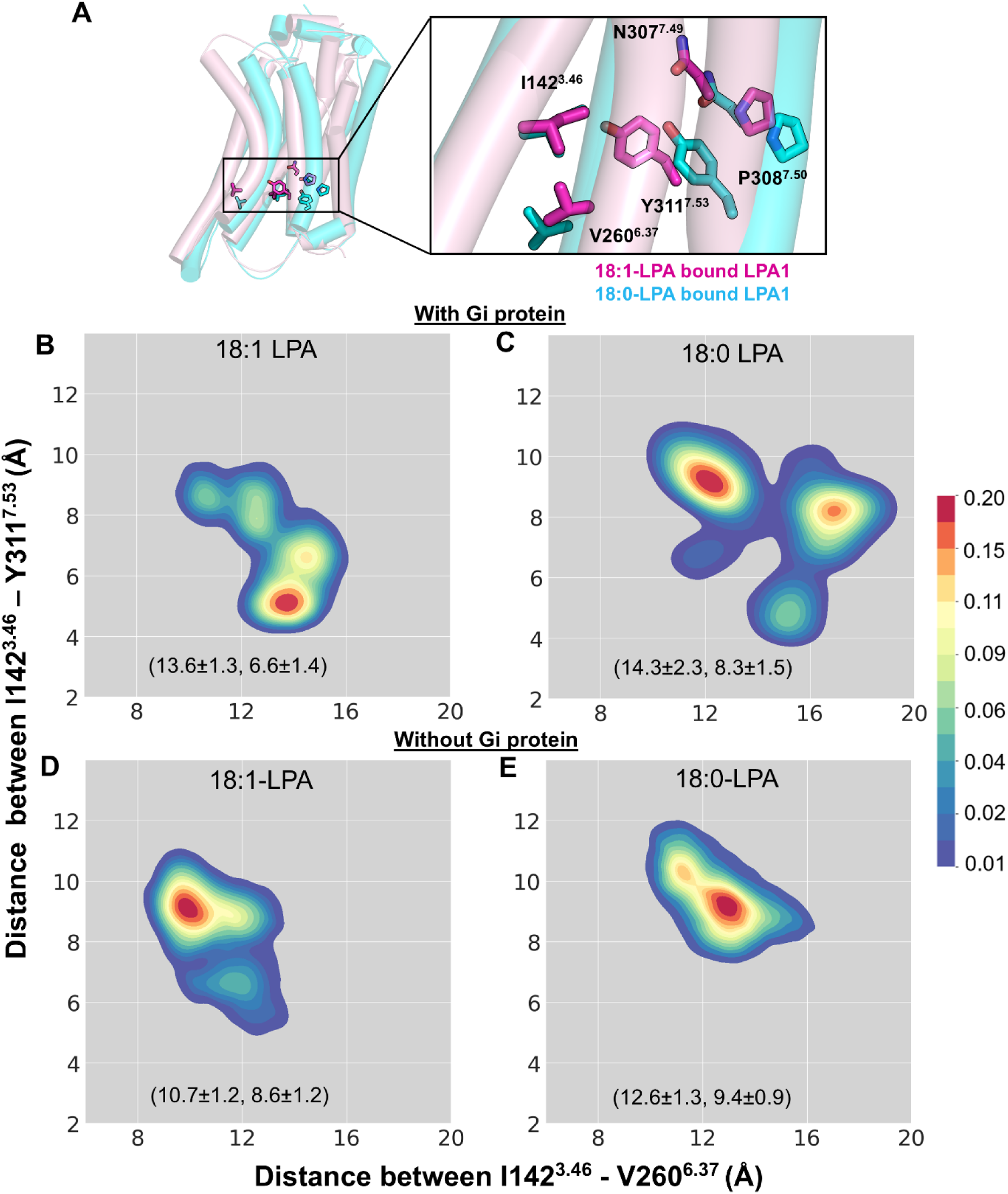
Differences in the interhelical activation-related distances between TM3-TM7 and TM3-TM6 of LPA1 provide the structural basis for full and partial agonist activity. (A) Critical residues of the activation switch, including the conserved NPxxY structural motif, TM3 (I142^3.46^), TM6 (V260^6.37^), and TM7 (Y311^7.53^), are shown from the superimposed cartoon representation of 18:0-LPA (cyan)- and 18:1-LPA (magenta)-bound LPA1 systems after 500 ns MD simulations. (B-E) 2D kernel density plots show the preferred TM3-TM6 and TM3-TM7 distances in four simulated systems. The closer interaction between TM3 (I142^3.46^) and TM7 (Y311^7.53^) is preferentially maintained in the 18:1-LPA-bound LPA1 with Gi (B) but not with the 18:0-LPA-LPA1-Gi system. The absence of Gi led to an increase in TM3-TM7 distance and a decrease in TM3-TM6 distance in both 18:1-LPA-LPA1 (C) and 18:0-LPA-LPA1 (D) receptor-only systems.

Overall, in the 18:1-LPA-LPA1 system in complex with the heterotrimeric Gi protein, the most predominant conformational change of the receptor involves a closer distance of about 4-5 Å between Y311^7.53^ and I142^3.36^ and a greater distance of approximately 13 Å between I142^3.36^ and V260^6.37^ **(Figure 4B)**. This implies that 18:1-LPA can restrict the conformation of the LPA1 receptor bound to the Gi protein in the active state for the cumulative simulation time of ∼1.5 microseconds. Interestingly, in the absence of Gi, there was an outward movement of TM7, leading to an increase in the TM3-TM7 distance by ∼4-5 Å and an inward movement of the TM6 by ∼4 Å **(Figure 4D)**. This difference is observed despite a similar binding pose and residue interactions of 18:1-LPA in the binding pocket of LPA1 with and without Gi protein.

With 18:0-LPA, LPA1 partially sampled a state with reduced TM3-TM7 distance (4-5 Å) and increased TM3-TM6 distance (15-16 Å) in the presence of heterotrimeric Gi protein. In the most dominant conformations, there was ∼9 Å distance between TM3-TM7 **(Figure 4C)**, which does not reflect a fully active state of LPA1. However, there was also an outward movement of the TM6 reflected by two clusters at ∼12 Å and ∼16 Å between TM3 and TM6; hence, the receptor is neither inactive. This observation likely reflects an intermediate state of the LPA1 receptor. Since 18:0-LPA is a partial agonist, it is unable to completely restrain the receptor in a fully active state despite the presence of the Gi protein **(Figure 4C)**. Without the heterotrimeric Gi protein, 18:0-LPA maintains the receptor in intermediate states similar to those observed with Gi-protein **(Figure 4E)**.

In addition to I142^3.46^-Y311^7.53^ distances, we analyzed ligand-induced differences in residue-residue distances at several other activation switches (**Figure S5A-C**). Importantly, in the presence of 18:1 LPA, V70^1.53^ shifts further away from Y311^7.53^ with an average distance of ∼10Å which closely resembles the measured distance in the active LPA1 (PDB ID: 7TD1) structure **(blue dotted line in Figure S5D)**. Interestingly, this distance was maintained in both the presence and absence of the heterotrimeric Gi protein. In contrast, the presence of 18:0 LPA resulted in a closer distance between V70^1.53^ and Y311^7.53^ that more closely resembles the measured distance in the inactive (PDB ID: 4Z34) structure **(red dotted line in Figure S5D)**. These 18:0 LPA-induced effects were also independent of the presence of the Gi protein. Analyses of distances between E145^3.49^ and R146^3.50^ and L256^6.33^-R146^3.50^ revealed similar patterns among the two LPA species in the presence and absence of Gi protein (**Figure S5E-F**).

### Aromatic residue conformations support ligand-induced activation states

One of the most commonly described structural motifs in class A GPCRs is the C^6.47^W^6.48^xP^6.50^ motif, which is located near the floor of the orthosteric binding site in LPA1^42^ (**Figure 5A**). In the inactive structure (PDB ID: 4Z34), L132^3.36^ forms a notable alkyl-π interaction with W271^6.48^ from the CWxP motif at a distance of ∼4.3 Å, indicated by a dotted line in **Figure 5B**. This interaction is absent in the active structure (PDB ID: 7TD1) in the presence of an agonist. This disruption appears to be partly due to the agonist-induced rotameric switching of neighboring W210^5.43^ at the base of the pocket. The chi2 dihedral angle of W210^5.43^ is 92 and -100 degrees in the active and inactive states, respectively **(Table S3)**. In our simulations, the average chi2 dihedral angles of W210^5.43^ in the presence of 18:0-LPA and 18:1-LPA are 80.6±11.8 and 92.9±6.6 degrees, respectively, which are near the active state conformation of LPA1 **(Figure 5C)**. 18:0-LPA and 18:1-LPA differ significantly in their ability to maintain other key activation switches, such as W271^6.48^, F267^6.44^, and Y311^7.53^ of conserved structural motifs, C^6.47^W^6.48^xP^6.50^, P^5.50^I^3.40^F^6.44^, and N^7.49^P^7.50^xxY^7.53^, in active state conformations. In the presence of 18:1-LPA, the chi2 dihedral angles of F267^6.44^ and W271^6.48^ are around -95° and 106°, respectively, and remain stabilized around the active state conformations **(Figure 5D-E & Table S3)**^25^. However, in the presence of 18:0-LPA, the mean dihedral angle of W271^6.48^ is around 27 ° and there is a limited sampling around the active state conformation **(Figure 5E)**. Also, the dihedral angle of F267^6.44^ shows wide variations (-3.9±71.9) with two clusters, likely representing intermediate states **(Figure 5D)**. In one of the clusters, the conformation of F267^6.44^ is synonymous with the active state, while in the other cluster, the F267^6.44^ dihedral is similar to the inactive state (**Table S3**).

**Figure 5.**
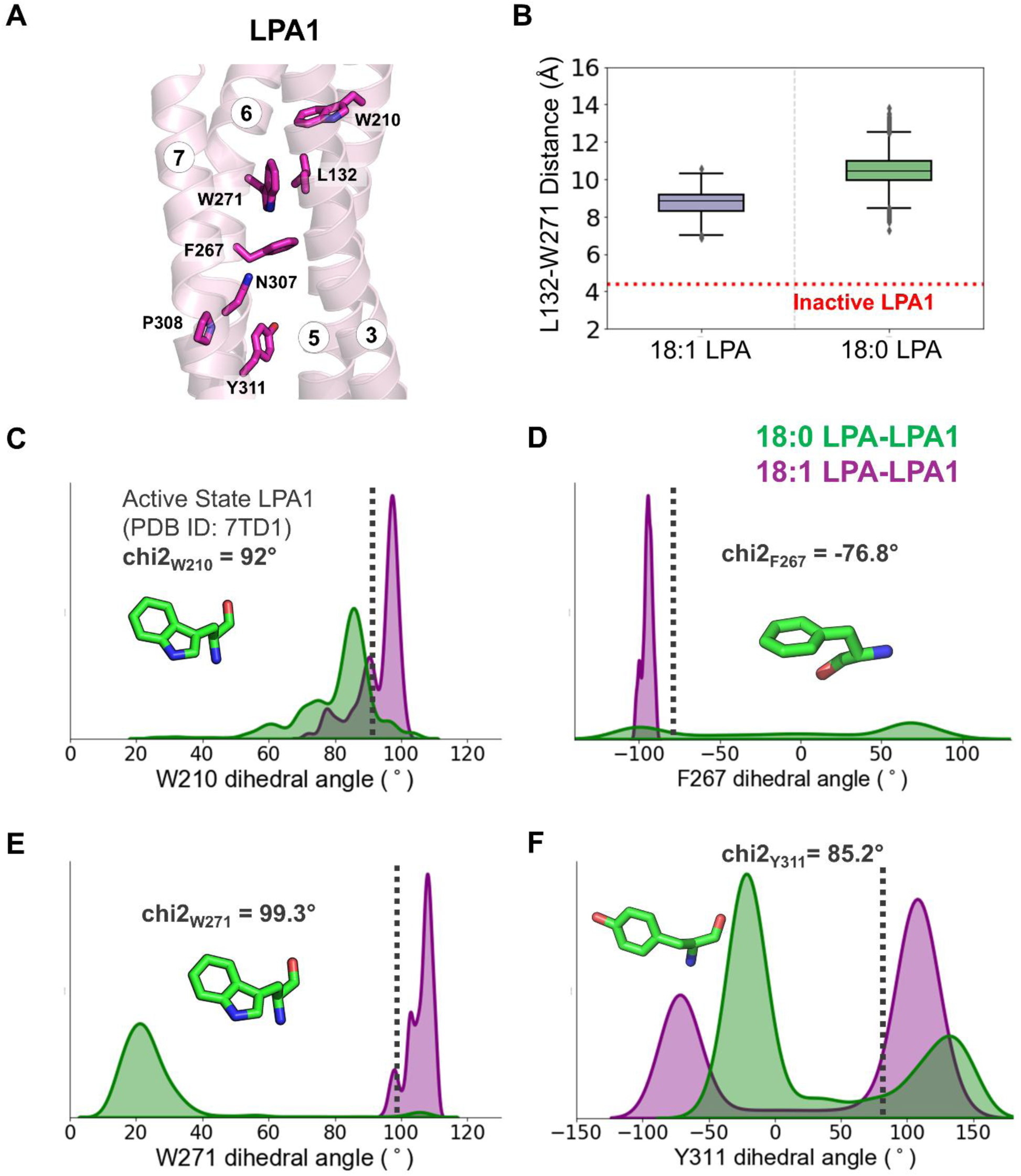
Ligand-induced conformational changes in residues that are part of conserved structural motifs at the base of the binding pocket. (A) Transmembrane helices 3, 5, 6, and 7 in a cartoon representation showing conserved residues forming specific structural motif interactions LPA1. (B) The box plot shows the disrupted alkyl-π interaction distance between residue W271^6.48^ and L132^3.36^ in the presence of 18:0- (green) and 18:1-LPA (purple). This interaction appears to be intact with a distance of around 4Å in the inactive state (PDB ID 4Z35), as indicated by the dotted red line. (C-F) Density plots show the chi2 dihedral angles for W210^5.43^, F267^6.44^, W271^6.48^ and Y311^7.53^ for both 18:0-LPA- (green) and 18:1-LPA-bound LPA1 (purple). In each plot, the dotted vertical line shows the corresponding value (also given as text) observed in the active state LPA1 (PDB ID 7TD1).

Interestingly, within the N^7.49^P^7.50^xxY^7.53^ motif, Y311^7.53^ underwent significant conformational changes in the presence of 18:0- and 18:1 LPA, resulting in a bimodal distribution of its dihedral angles in both systems. However, the distribution toward the active state is more prominent in the presence of 18:1-LPA than in the presence of 18:0-LPA (**Figure 5F**). In the case of P308^7.50^, both 18:0- and 18:1-LPA allowed the receptor to sample the dihedral angles mostly near the active state, although the angles appeared in the inactive state to some extent **(Figure S6)**. The dihedral changes and displacement noted for the C^6.47^W^6.48^xP^6.50^, P^5.50^I^3.40^F^6.44^, and N^7.49^P^7.50^xxY^7.53^ motifs play a crucial role in creating an interface for receptor Gi protein interaction. Hence, differential modulation of these motifs in the presence of 18:0- and 18:1-LPA could play a part in their differential activities at the receptor and the consequent downstream responses, such as ERK activation.

### Ligand-induced conformational changes affect LPA1 and Gi protein interactions

In GPCRs, ligand-induced receptor activation signals are transmitted through a series of activation switches, eventually resulting in changes at the intracellular interface that facilitate and maintain G-protein coupling and interactions **(Figures 6A and S7A)**. Our MD simulation results are in agreement with the recent cryo-EM study (PDB ID: 7TD1) that the receptor interface in LPA1 includes the intracellular ends of TM3-TM6 as well as the various intracellular loops, especially ICL2 **(Figures 6A and S7A)**^25, 26^. These regions are involved in very extensive polar interactions, particularly salt bridges with the C-terminal end of the G_αi_-α5 helix of the G-protein.

**Figure 6.**
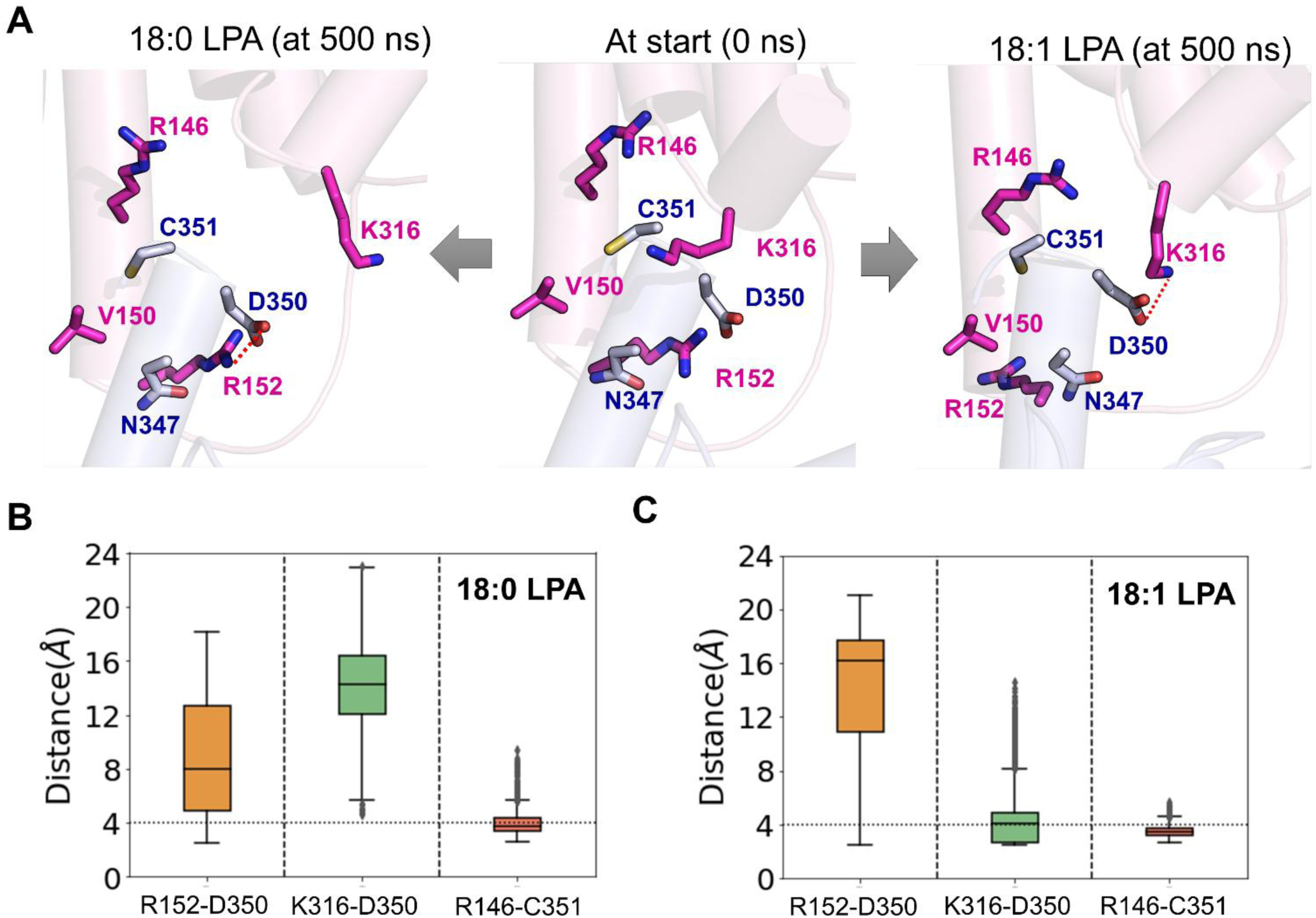
18:0- and 18:1-LPA distinctly affect LPA1-Gi protein interactions. (A) Dynamics and critical residue interactions at the LPA1-Gi protein interface in the presence of two LPA species at the beginning (time = 0 ns) and end of 500 ns MD simulations. LPA1 and Gi protein are shown in cartoon representations in pink and light blue; critical residues of the respective interacting species are shown in licorice in magenta and blue, respectively. (B and C) Boxplots of distances between critical residues forming important polar interactions at the interface between LPA1 and Gα-helix of the G-protein in 18:0-LPA and 18:1-LPA bound systems, respectively. The strong salt bridge between K316^H8^ and D350^G.H5.22^ observed in the presence of 18:1-LPA is absent in the 18:1-LPA system.

One of the important determinants of the receptor-Gi-protein coupling is the hydrogen bond between R146^3.50^ (from the E^3.49^R^3.50^H^3.51^ motif) and the backbone carbonyl atom of C351^G.H5.23^ from the Gα-helix. In both the 18:0- and 18:1-LPA-LPA1 systems **(Figure 6B-C)**, this interaction was maintained entirely during the 500 ns simulations **(Figure 6C-D)**. In addition, C351^G.H5.23^ engages in polar interactions with T149^3.53^ and V150^3.54^ residues at the intracellular ends of TM3. These interaction profiles were similar with both ligands **(Figure S7B-C)**.

Not surprisingly, the differential movement of the TM6 and TM7 helices, as outlined earlier, could result in alterations at the receptor-Gi-protein interface **(Figure 4A-E)**. For example, the cryo-EM structure of LPA1-LPA (PDB ID:7TD1) features a salt bridge between R152^3.56^ from the receptor and D350^G.H5.22^ of the G-protein. This interaction was disrupted to a much greater extent in the 18:1 simulation than in the 18:0-LPA -LPA1 system **(Figure 6C-D)**. The differential movement of R152^3.56^ resulted in differential contact with I343^G.H5.15^ of the Gα helix in the 18:0-LPA bound systems **(Figure S7B-C)**. However, the strong polar interaction of R152^3.56^ with N347^G.H5.19^ was well preserved in both systems.

In the 18:1-LPA-LPA1 simulations, the inward movement of TM7 is sustained, which led to the maintenance of a very stable interaction between K316^H8^ at the intracellular end of the receptor and D350^G.H5.22^ in the Gα-helix. This interaction was noticeably absent in 18:0-LPA-LPA1 simulations, which might be unrelated to the outward movement of TM7 **(Figure 6B-C)**. D350^G.H5.22^ (Gα-helix) contacts R146^3.50^ to a greater extent in the 18:1-LPA-LPA1 simulations than in the 18:0-LPA simulations **(Figure S7B-C)**.

The intracellular loops of GPCRs have been shown to participate to varying extents in G-protein coupling. Differential interaction with the ICL3, which occurs with the S1P1 receptor, was previously proposed as a difference with the LPA1 interface, where these interactions are less significant. In our simulations with 18:0- and 18:1-LPA, the ICL2 residues, including M153^ICL2^ and Q154^ICL2^, are involved in hydrophobic interactions with I343^G.H5.15^ for the majority of the simulation time. In 18:1-LPA simulations, Q154^ICL2^ also engages in significant polar interactions with N347^G.H5.19^ that were less prevalent in 18:0-LPA simulations **(Figure S7B-C)**. In addition, hydrophobic interactions between L256^6.33^ in TM6 and L353^G.H5.25^ of the Gα-helix were observed as common to both systems. TM5 residues, including R235^5.68^, R238^5.71^, and M239^5.72^, also play a role in G-protein coupling via polar and hydrophobic interactions with F334^G.H5.06^, D337^G.H5.09^, and D341^G.H5.13^ of Gi protein **(Figure S7B-C)**.

### Formation of water channels across the transmembrane helices

Recent studies have shown that water networks could modulate signal transduction from the extracellular region to the intracellular loop regions of GPCRs. Specifically, the presence of large water clusters near the intracellular region of TM7 N^7.49^P^7.50^xxY^7.53^ motif has been observed in mu-opioid, β-adrenergic, and adenosine receptors^43–45^. In our LPA1 simulations with both 18:0- and 18:1-LPA, we observed the gradual expansion of continuous water channels that run from the extracellular to intracellular ends of the receptor **(Figure 7A-B)**. Particularly, there was a bulge in the channel at the intracellular end corresponding to the region of the N^7.49^P^7.50^xxY^7.53^ motif. This differs from the inactive LPA1 structure that has been reported to have a discontinuous channel, likely due to water occlusion in the TMD areas^46^. To confirm the likelihood that this channel conducts water molecules, we measured water flux between the upper and lower ends of the binding pocket **(Figure 7E-F)**. In both systems, at least one water molecule moves between the upper and lower regions of the receptor for a significant period during the simulation time. Water flow through the channel was slightly greater in the 18:0-LPA system compared to 18:1-LPA; this is consistent with the observation that the 18:0-LPA-LPA1 system features a more extensive water channel **(Figure 7A-B)** between the intracellular and extracellular regions.

**Figure 7.**
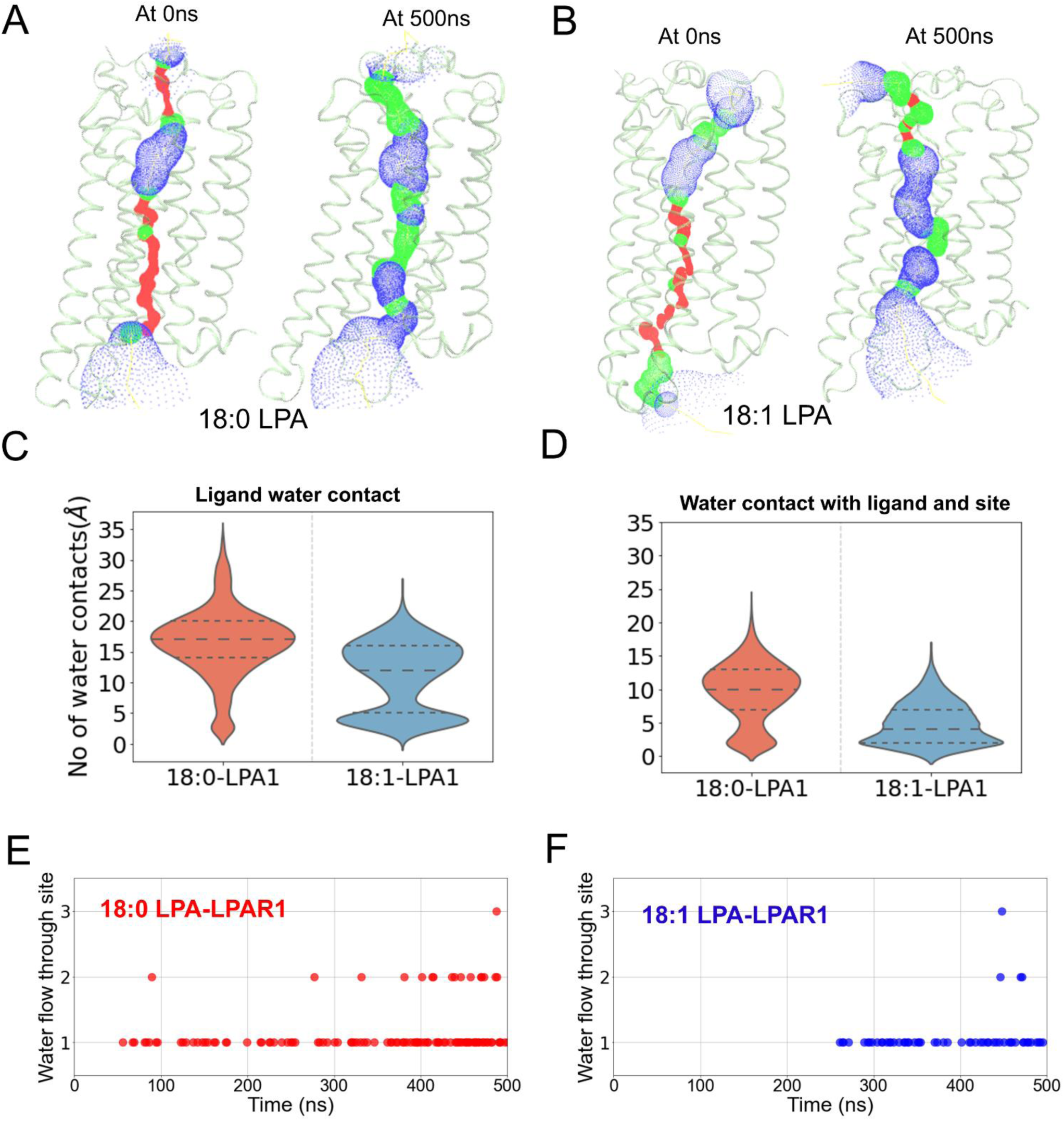
**18:0- and 18:1-LPA induce the formation of water channels in LPA1**. (A-B) Opening of an intracellular water channel along the transmembrane helices during the simulation in 18:0-LPA (A) and 18:1-LPA bound LPA1-Gi-protein systems. (C) Ligand-water contacts within the receptor. This was measured by computing the total number of water molecules within 4 Å of 18:0- (red) and 18:1-LPA (blue) species. (D) Water molecules that are in contact with both the binding pocket residues and the ligand. (E-F) The number of water molecules that flow from the extracellular part of the receptor through the base of the binding pocket in 18:0-LPA (E) and 18:1-LPA (F) bound LPA1.

In addition to the water channel through the orthosteric binding site, we also observed a significant amount of ligand water contact that was generally higher for 18:0-LPA **(Figure 7C)**. At the start of the simulation, there was more water contact with the glycerol-phosphate groups and little to no water contact with the alkyl chains **(Figure S8A-B)**. Alkyl chain water contact emerged with the expansion of the water network passing through the binding site. Also, we observed several mobile water molecules in the orthosteric site in the presence of both 18:0 and 18:1 LPAs. **(Figure 7D)**. However, no specific lasting water-mediated ligand interactions were observed.

### Optimal communication paths between the activation site and G-protein coupling interface

To obtain insights into the signal transduction pathway between orthosteric site residues and residues that interface with Gi-protein at the intracellular side, we carried out dynamic network analyses using the VMD plugin Network View. The optimal residue-to-residue communication paths between crucial binding site residues (R124^3.28^, W271^6.48^, and K294^7.36^) and residues that engage with the Gi-protein interface (R146^3.50^, R152^3.56,^ and K316^H8^) were analyzed based on the MD trajectory data.

In the 18:1-LPA bound LPA1-Gi protein system, a single optimal path connected R124^3.28^ in TM3 with R146^3.50^ and R152^3.56^, which directly interact with the Gα helix of the Gi protein **(Figure 8A)**. This path spans only the TM3 helix and includes I142^3.46^, which was noted earlier as having closer contact with Y311^7.53^ in the 18:1-LPA-LPA1 system compared to the 18:0-LPA-LPA1 system **(Figure 4A-C)**. However, in the 18:0-LPA-LPA1 system, two optimal paths connect R124^3.28^ simultaneously to R146^3.50^ and R152^3.56^. The path running from R124^3.28^-R146^3.50^, including residues (L127^3.31^, S131^3.35^, and S135^3.39^), resembles that of the 18:1-LPA-LPA1 receptor system more closely than the R124^3.28^-R152^3.56^ path.

**Figure 8.**
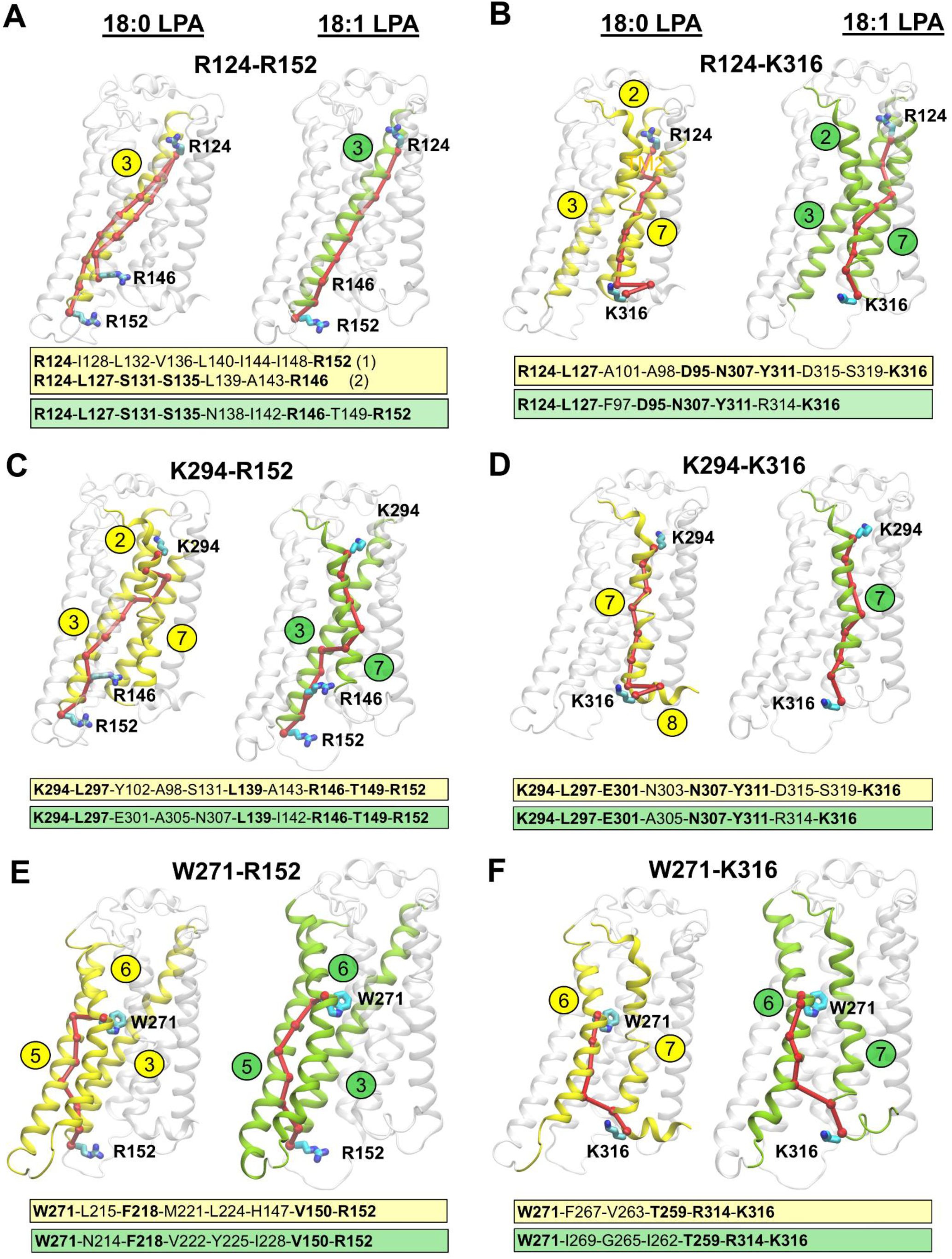
Optimal signal transduction paths in 18:0-LPA- and 18:1-LPA- bound LPA1 systems, as determined by network analyses. The optimal paths were calculated between critical binding site residues and residues at the Gi-protein coupling interface. In each case, 18:0-LPA- and 18:1-LPA-activated LPA1 receptors are depicted in cartoon representations and participating helices are colored and labeled in yellow and green, respectively. A-B) The optimal paths between the orthosteric site residue R124^3.28^ and residues at the Gi protein-coupling interface R152^3.56^ and K316^H8^, respectively. C-D) The optimal paths between K294^7.36^ with R152^3.56^ and K316^H8^, respectively. E-F) The optimal paths between W271^7.36^ and R152^3.56^ and K316^H8^, respectively. For each path, all the residues along the entire path for 18:0- and 18:1-LPA bound LPA1 are given in yellow and green text boxes, respectively. Common residues among the paths are in bold fonts.

The optimal communication path between R124^3.28^ and K316^H8^ passes along TM2, TM3, and TM7 helices in both 18:0- and 18:1-LPA bound systems. One common residue along this path, D95^2.50^, is conserved across all LPA (1-6) receptors **(Figure 8B and Figure S11)**. Interestingly, this path also included N307^7.49^ and Y311^7.53^ from the N^7.49^P^7.50^xxY^7.53^ motif in both systems with 18:0 and 18:1-LPAs. The noticeable difference here is the presence of R314^7.56^ in the 18:1-LPA1 system, unlike the 18:0-LPA1 system, which includes D315^H8^ and S319^H8^ from the helix 8.

Also, the residue communication path between K294^7.36^ and R152^3.56^ transverses TM3 and TM7 in the 18:1-LPA1 system. Similar to the R124^3.28^-R152^3.56^ path, it included residue I142 from TM3. The path also features N307^7.49^ from the N^7.49^P^7.50^xxY^7.53^ motif. In contrast, this same path (K294^7.36^-R152^3.56^) in the 18:0LPA liganded system also transverses TM2 (A98^2.53^ and Y102^2.57^) and did not pass through the I142^3.46^ residue. Across all K294^7.36^ paths with the G-protein interface residues (R152^3.56^ and K316^H8^), L297^7.39^, located near the bottom of the binding site, is the most commonly occurring residue **(Figure 8C&8D)**.

Unlike the earlier connections analyzed, the optimal residue communication path between W271^6.48^-R152^3.56^ passes through the TM5 in the presence of both ligands **(Figure 8E&8F)**. The common residue here, F218^5.51^, is conserved across the EDG LPA (1-3) receptors. This path also included V150^3.54^, which interacts with the Gi protein in both 18:0-LPA1 and 18:1-LPA receptor systems. Also, the W271^6.48^-K316^H8^ path noticeably did not pass through the N^7.49^P^7.50^xxY^7.53^ motif with either 18:0 or 18:1-LPA ligands bound to the LPA1 receptor. The only common residues with both ligands across this path include T259^6.36^ and R314^7.56^. The uniqueness of the W271^6.48^ to G-protein interface communication path prompted us to evaluate its optimal connection pathway with site residues (R124^3.28^, K294^7.36^) and the N^7.49^P^7.50^xxY^7.53^ motif. Consequently, we observed significantly different connections of W271 to (R124^3.28^, K294^7.36^, and Y311^7.53^) between 18:0-LPA and 18:1-LPA bound LPA1 receptor systems **(Figure S9A-C)**.

### Access and binding mechanisms of 18:0 and 18:1-LPAs to LPA1

To investigate the ligand access routes and binding to the orthosteric binding pocket, we implemented association simulations using well-tempered metadynamics (WT-metaD), one of the most efficient enhanced sampling techniques. We performed these simulations in multiple replicates to characterize the plausible access routes for 18:0-LPA into the receptor. The free-energy surface for the binding process was characterized using two collective variables: 1) the distance between the center-of-mass (COM) of the ligand and COM of the binding site residues and 2) the internal angle of the ligand accounting for its conformational changes during the binding process. All association simulations started with ligands in the extracellular aqueous phase at ∼20 Å away from the binding site residues **(Figure 9A)**.

**Figure 9.**
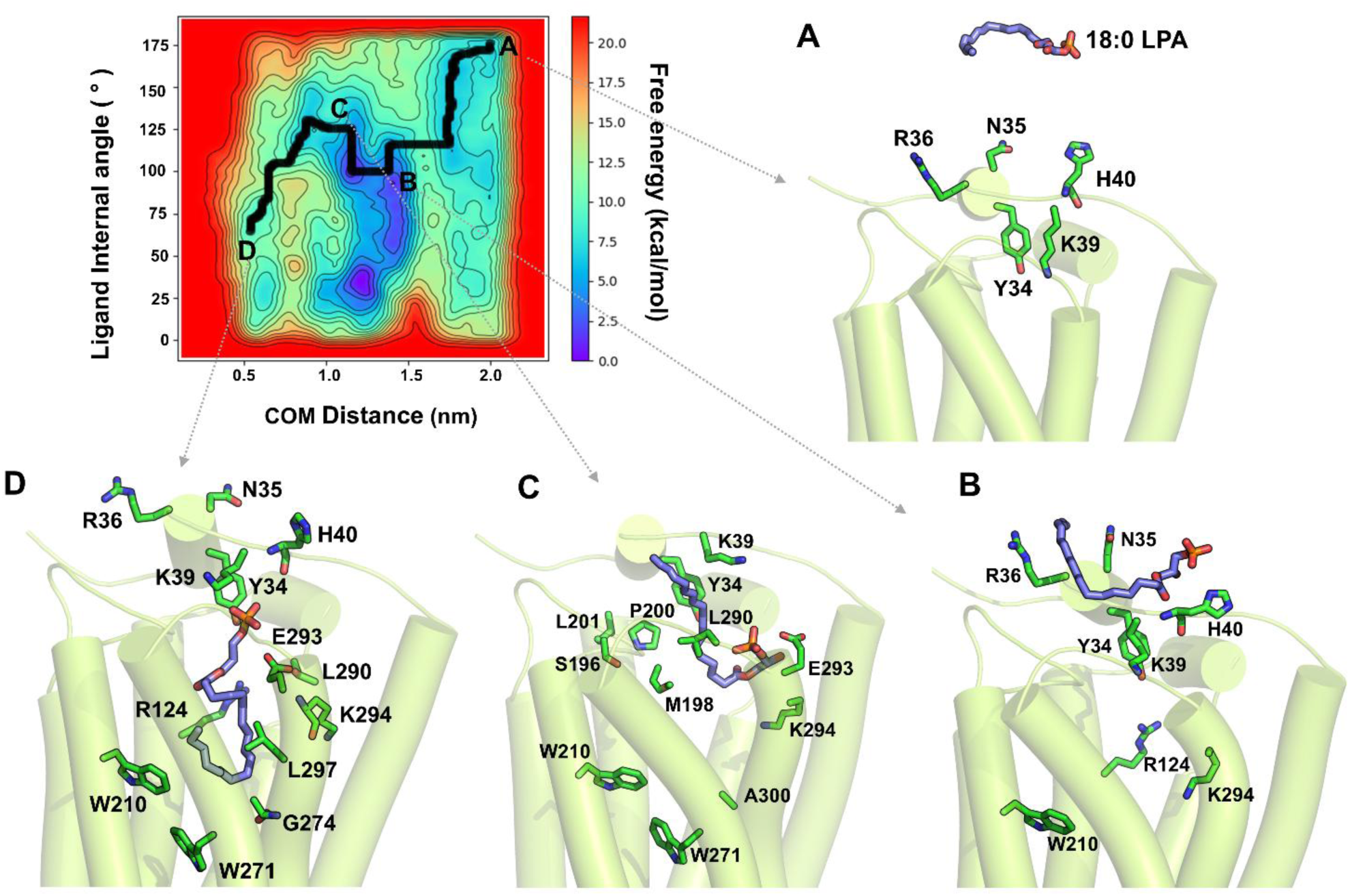
The free-energy surface (FES) of 18:0-LPA’s access and binding to the LPA1 receptor from the aqueous phase. The 2D FES is depicted using two collective variables: 1) the center-of-mass distance between the ligand and binding site residues (x-axis) and 2) the ligand’s internal angle calculated between its phosphate (P1) head, middle (C9-C10), and lower alkyl chain carbons (C1-C2). The minimum energy path of the ligand to the receptor, as determined by WT-metaD, is depicted in the bold black line. (A-D) Representative starting, intermediate, and bound conformations are shown. (A) 18:0-LPA is initially located ∼20 Å away from the receptor in the aqueous phase. (B) 18:0-LPA rapidly approaches the receptor’s N-terminal loop; the phosphate head group of the ligand contacts K39 of the N-terminal first and forms polar interactions with K39^Nterm^, H40^Nterm^, Y34^Nterm^, and R36^Nterm^. (C) The 18:0-LPA alkyl chain undergoes significant conformational changes and attempts to enter the receptor via the space between the ECL1 and ECL2 loops. (D) The pouch-like shape of the binding pocket forces the 18:0-LPA alkyl tail to bend, forming hydrophobic contacts with R124^3.28^, Q125^3.29^, G274^6.51^, E293^7.35^, and K294^7.36^ in TM3 while maintaining polar contacts with N-terminal Y34^Nterm^.

Often, 18:0-LPA rapidly approached the membrane with its polar glycerol-phosphate head group engaging with the lipid headgroups. Initially, the ligand underwent extensive conformational changes, establishing contacts with the N-terminal loop residues through the polar groups. Specifically, the ligand interacted with Y34^Nterm^, N35^Nterm^, R36^Nterm^, and K39^Nterm^ for nearly 40% of the simulation time (∼15 ns) before it entered the orthosteric pocket **(Figure 9B)**. With the polar headgroups engaged in these interactions, the alkyl chain curled to reduce its surface area exposed to water.

As 18:0-LPA approached the pocket entrance, it underwent a rapid and significant conformational change from an extended orientation, approximately parallel to the membrane plane, to a nearly perpendicular acute orientation. This is further evident from the ligand’s internal angle, which changed from ∼160° to ∼60°, as observed in the thermodynamic free-energy diagram **(Figure 9C)**. Beyond the N-terminal, the alkyl chain first contacted residues P200^ECL2^ and M198^ECL2^, as it gained access to the binding pocket and subsequently contacted S196^ECL2^ and L201^ECL2^ **(Fig.9C)**. The molecule entered through the space between the ECL loops, and closer to ECL2. Within the transmembrane region, it made first contact with TM7 residues, L290^7.32^ and E293^7.35^ of the binding site. Further, the ligand moved deeper into the pocket, with its alkyl chain establishing contacts with residues R124^3.28^, G274^6.51^, K294^7.36^, and L297^7.39^ from TM3, TM6, and TM7 helices. Subsequently, 18:0-LPA assumed a bent conformation with the alkyl chain pushing against TM7 **(Figure 9D)**. In its final bound orientation, the phosphate headgroup formed polar contacts with E293^7.35^ while maintaining a hydrogen bond interaction with Y34^Nterm^ **(Figure 9D)**.

18:1-LPA first approached the N-terminal loop from ∼20 Å in the aqueous phase in a manner similar to the 18:0-LPA entry **(Figure 10A)**. The phosphate headgroup of the ligand engaged in extensive polar interaction with N-terminal residues, such as Y34^Nterm^, N35^Nterm^, R36^Nterm^, and K39^Nterm^, first and subsequently with R36^Nterm^, Y34^Nterm^, and sparsely with H40^Nterm^. These N-terminal interactions were sustained throughout most simulation time **(Figure 10B)**.

**Figure 10.**
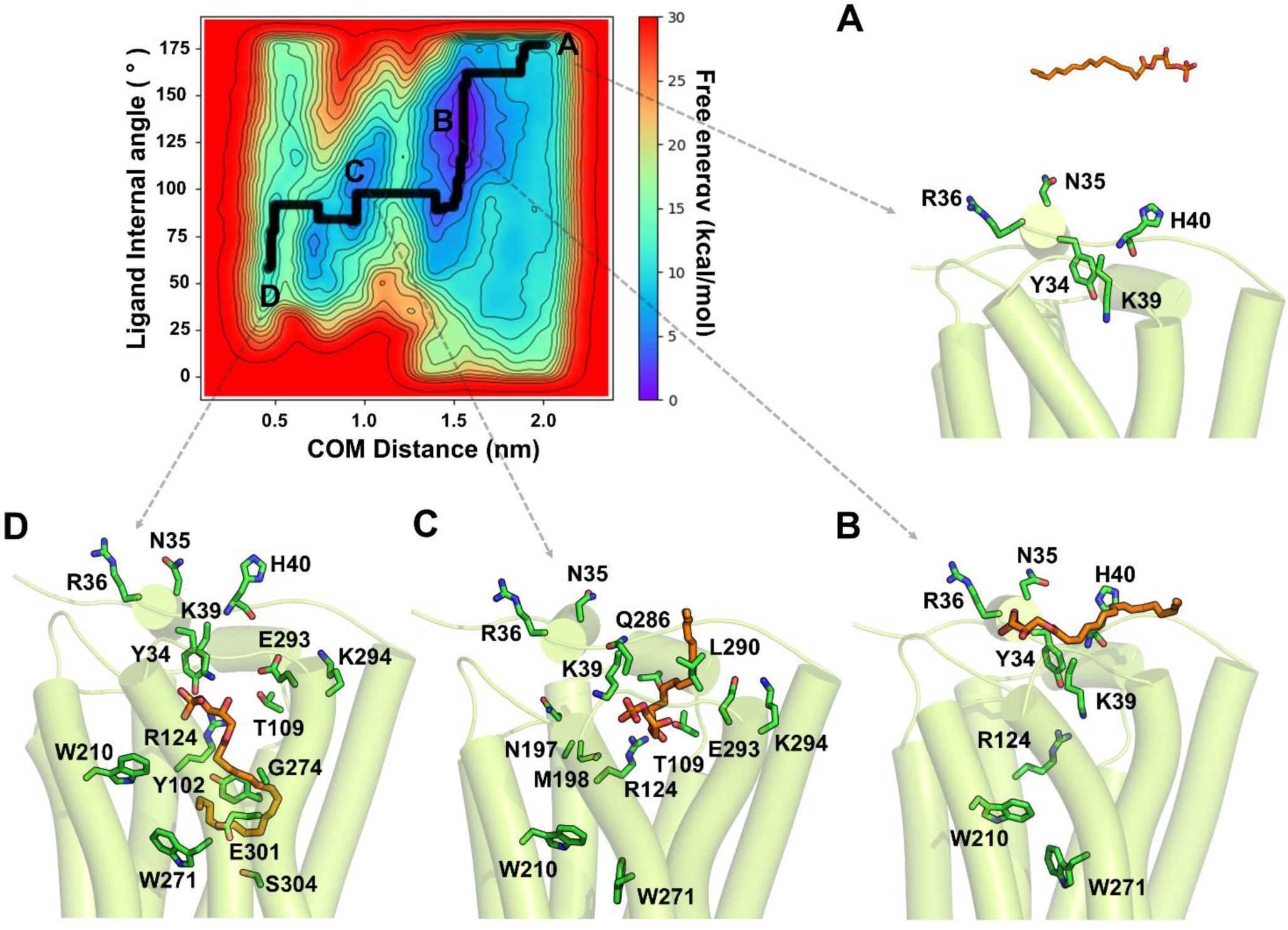
The free-energy surface (FES) of 18:1-LPA’s access and binding to LPA1. The 2D FES is depicted using two collective variables: the distance between the center-of-mass of the ligand and that of binding site residues (x-axis) and the ligand’s internal angle. The minimum energy path of the ligand to the receptor is represented by the bold black line, and major intermediate conformations are labeled A-D. (A) 18:1-LPA is initially located within the aqueous bulk at a distance of ∼20 Å from the receptor. (B) 18:1-LPA rapidly approaches the N-terminal loop of the receptor. While the phosphate head group first contacts K39^Nterm^ in the N-terminal and forms polar interactions with K39^Nterm^, H40^Nterm^, and R36^Nterm^, the alkyl chain partitions into the membrane. (C) 18:1-LPA enters the receptor via the extracellular loops. (D) As 18:1-LPA approaches the binding site, its lower alkyl chain comes in contact first with L290^7.32^ and E293^7.35^. The ligand adopts a final pose with a bent alkyl conformation.

Similar to 18:0-LPA, we observed that 18:1-LPA entered the receptor from the spaces between the extracellular loops. The ligand made contact with P200^ECL2^, M198^ECL2^, N197^ECL2^, T109^ECL1^, and Q286^ECL3^ upon entering the binding site **(Figure 10C)**. Subsequently, it entered the binding region between the TM helices while establishing contacts with L290^7.32^, E293^7.35^, and K294^7.36^ along the path. The alkyl chain of 18:1-LPA also underwent a significant conformational change with an eventual bent conformation on entry. This is evident in the sharp decrease of the ligand’s internal angle from ∼ 150° to ∼ 60° on the free-energy surface diagram **(Figure 10B-C)**. In the final bound pose, the alkyl chain established hydrophobic contacts with Y102^2.57^, R124^3.28^, W210^5.43^, W271^6.48^, G274^6.51^, E301^7.43^, and S304^7.46^, which are all part of the receptor’s orthosteric pocket **(Figure 10D)**. The ligand also sustained the polar interactions with Y34^Nterm^ and K39^Nterm^ of the N-terminal and formed a new hydrogen bond with T109^ECL1^.

### Membrane Partitioning Characteristics of 18:0 and 18:1-LPA Species

Previous studies from our lab indicated that despite having similar lipophilic-amphiphilic characteristics, ligands can exhibit distinct membrane partitioning characteristics and take separate paths to reach the orthosteric sites in class A GPCRs^47^. Therefore, to elucidate the likely differences in membrane partitioning characteristics between 18:0-LPA and 18:1-LPA, we used steered molecular dynamics (SMD) and umbrella sampling (US) simulations of the ligands in a model membrane consisting of 1-palmitoyl-2-oleyl-sn-glycero-3-phosphocholine (POPC) and cholesterol. The potential of mean force (PMF) curve **(Figure 11B)** derived from the US simulations revealed the free energy of solvation for both ligands partitioning from water into the membrane along the bilayer normal (z-axis). The free-energy minima and maxima on the PMF curve indicate the most and least energetically favorable locations for the ligands. Although both ligands show favorable partitioning energy from water to the membrane, the phase transfer process for 18:0-LPA (ΔG_partitioning_ = -9.3 ± 0.05 kcal/mol) appeared to be more energetically favorable than 18:1-LPA (ΔG_partitioning_ = -7.2 ± 0.07 kcal/mol). Interestingly, the free-energy minima of the center-of-mass of both ligands **(Table S4)** are located at |Z _min_| ∼ 15 Å. Both ligands encountered an energetic barrier as they approached the membrane core. However, the energy barrier is relatively higher for 18:1-LPA (ΔG_crossing_ = 12.4 ± 0.08 kcal/mol) as compared to 18:0-LPA (ΔG_crossing_ = 11.1 ± 0.06 kcal/mol). For both ligands, the thermally accessible regions (RT =0.616 kcal/mol, temperature=310 K) extended for ≈ 2Å on either side of Z_min_.

**Figure 11.**
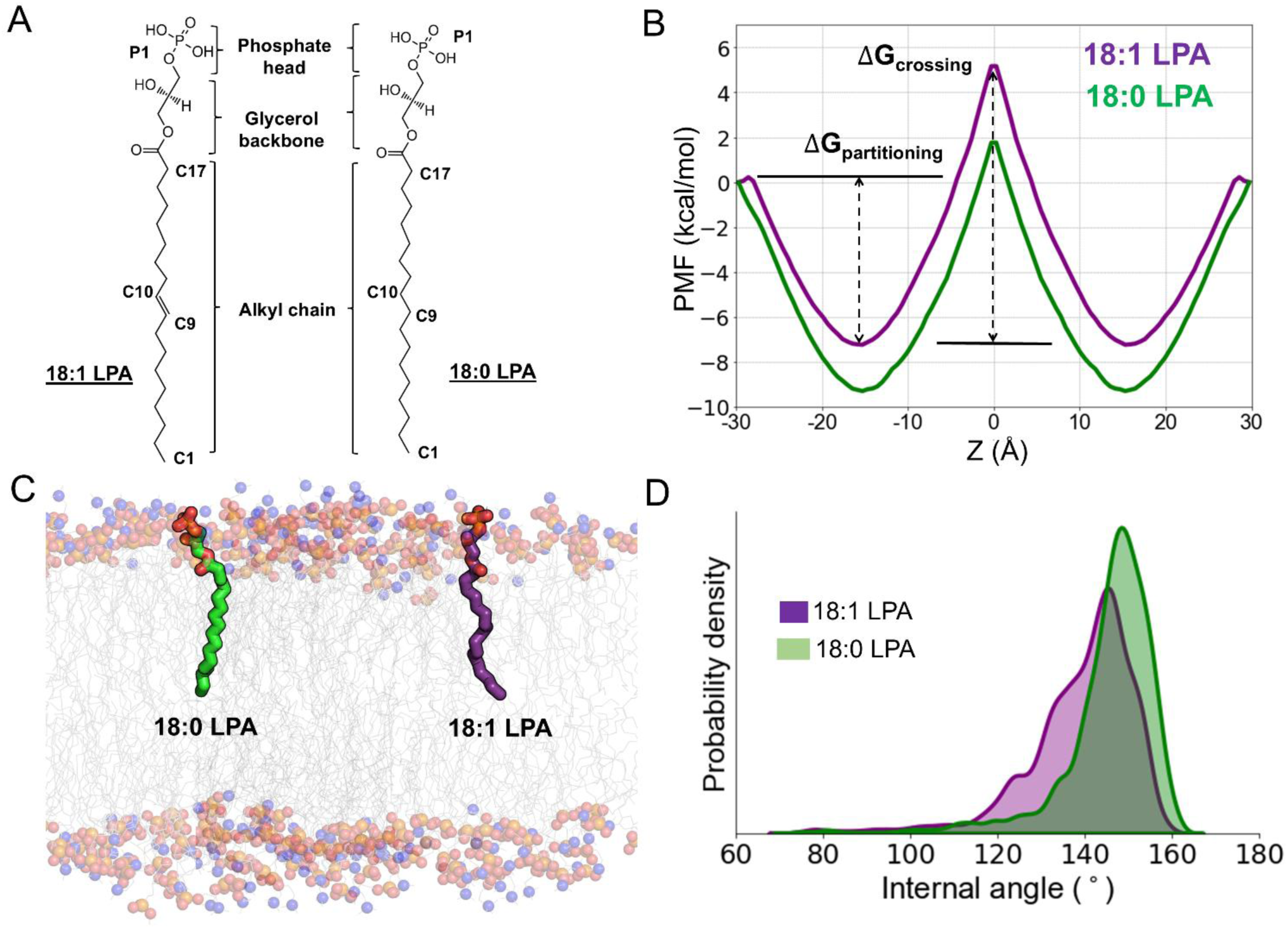
Membrane partitioning characteristics of 18:0 and 18:1-LPA species. (A) 2D-structures of the studied LPAs with their different functional groups and parts labeled. (B) The potential of mean force (PMF) curves for the ligands depicting the solvation-free energies for membrane partitioning and crossing the bilayer and energetically favorable bilayer locations. (C) The time-average preferred orientations of the ligands (in licorice representation) within the membrane. The membrane lipid head group atoms are depicted as balls: choline nitrogen (blue), glyceryl oxygen (red), and phosphorus of the phosphate group (olive green). The alkyl tails of the lipids are represented as lines (gray). (D) The probability density plot shows the internal angle of the ligands. The internal angle was measured between the phosphate head (P1), middle (C9-C10), and alkyl tail end (C1-C2) of the ligand.

To obtain further insights into the differences observed in the solvation-free-energy profile of both ligands, we analyzed the internal angle of the ligand within the bilayer. The angle was measured using three points representing the head, middle, and tail parts of the ligand (defined by the P1 group and the middle C9 and C1 carbon atoms of the alkyl chain). Not surprisingly, both ligands adopt a nearly extended conformation. The average conformation of 18:0-LPA (145.4° ± 11.2°) is such that the head tilts acutely towards the membrane bilayer normal while the alkyl chain adopts a nearly straight orientation in the membrane core to maximize its hydrophobic interactions with the lipid tails **(Figure 11C-D)**. The internal angle of 18:1-LPA is slightly lower (139.1° ± 12.5°) than 18:0-LPA. The probability distribution plot **(Figure 11D)** also showed that 18:1-LPA explored more conformations in the membrane than 18:0-LPA. This is evident in the relatively higher standard deviation.

Also, we calculated the atomic contacts (as % contact frequency), which indicates the fraction of the simulation time during which different parts of 18:0-LPA and 18:1-LPA are within 4 Å of the various functional groups of the lipids. These analyses were done using the simulation trajectory representing the low-energy windows |Z_min_| ∼ 15 ± 2 Å for both ligands **(Figure S10)**. In both ligands, the phosphate headgroup maintained strong contact with the choline functional groups of the lipids while staying further away from the glyceryl carbonyl and alkyl lipid groups. However, there was a slightly higher frequency in 18:1-LPA’s phosphate group contact with the choline lipid group. The glycerol backbone in both ligands had a high frequency of contact with all the lipid headgroups (choline, phosphate, glyceryl carbonyl). Again, the glycerol backbone of 18:1-LPA had a slightly greater contact than 18:0-LPA. Interestingly, a similar observation was noted in the ligand’s alkyl chain interaction, where both the upper and lower parts of the alkyl chain of 18:1-LPA had a higher frequency of contact with the lipid’s alkyl group than the alkyl chain of 18:0-LPA. Overall, 18:0 LPA and 18:1 LPA have very similar membrane partitioning profiles. Although the free energy of partitioning seems slightly more favorable for 18:0-LPA, 18:1-LPA explored more conformations and contacts with the membrane functional groups.

## DISCUSSION

The LPA1 receptor is implicated in the pathogenesis of various diseases, including neuropsychiatric disorders^48^, neuropathic pain^49^, angiogenesis^50^, and cancer^51–53^. Notably, the upregulation of LPA1 has been demonstrated in many cancers, including breast, prostate, pancreatic, lung, and hepatocellular carcinoma, in both cell culture and animal studies^54–57^. These studies consistently show that LPA1 is involved in tumor cell growth, proliferation, migration, and propagation of the tumor microenvironment. Therefore, antagonizing LPA1 to prevent its downstream signaling represents an attractive therapeutic strategy for treating cancer^58^ and other diseases, including pulmonary fibrosis ^59^ and cutaneous systemic sclerosis ^60^. Despite its role in multiple diseases, there are currently no clinically approved LPA1 modulators. Challenges include the cross-reactivity of LPA1 with other lysophosphatidic acid receptors (LPA2-6) due to their high sequence homology **(Figure S11)** as well as with other related GPCRs, including sphingosine-1-phosphate and cannabinoid receptors. In addition, LPA1 demonstrates notable promiscuity in G-protein coupling, binding to the alpha-subunits of heterotrimeric G_i/o_, G_12/13_, and G_q/11_, potentially modulating a wide variety of signaling pathways with similar or contradictory physiological effects^5^. Moreover, the receptor is activated by a wide variety of endogenous agonists, many of which are ubiquitous and possess varying efficacy^5^. Factors influencing the activity of similar endogenous agonists include the chain length and degree of unsaturation^10,23^.

This study shows that a double bond (unsaturation) in 18:1-LPA plays a role in the differential activation of LPA1 compared to 18:0-LPA in prostate cancer PC-3 cell lines. Immunoblotting assay results indicate that 18:1-LPA elicits greater Erk activation than 18:0-LPA at the same concentration (**Figure 1**). These efficacy differences suggest the possible influence of unsaturation in ligand interactions at the receptor’s binding site^15^. A thorough understanding of the differences in the receptor activation by these two LPA species may enable effective strategies for designing LPA1 selective modulators in different therapeutic areas. While most GPCRs share similar activation mechanisms, there are also unique differences noted among them^28^. Thus, elucidating unique activation mechanisms and examining how the activation signal is transmitted from the binding site residues to various activation switches along the transmembrane region is critical to a complete understanding of the activation process in LPA1.

Results from unbiased MD simulations of LPA1-LPA complexes show that both ligands exhibit reasonable stability at the binding site **(Figure 2A-D)**. The glycerophosphate head, identical in both ligands, has a similar pattern of contact and polar interactions with a few exceptions, such as K39^Nterm^ and T113^ECL1^, which interact more with 18:0-LPA. These differences are reflected in the binding energies, with K39^Nterm^ and T113^ECL1^ contributing more to the binding affinity of 18:0-LPA compared to 18:1-LPA. In addition, MM/PBSA calculations support the importance of the N-terminal loop residues, K39^Nterm^ and Y34^Nterm^, to ligand affinity. Also, the calculated energies reveal that R124^3.28^ and K294^7.36^ make the highest contributions to binding affinity. In contrast, residues lining the base of the pocket have limited contributions to affinity but seem to play a significant role in the differential receptor activation by the ligands.

Analyses of transmembrane helical distances and dynamics of conserved structural motifs offer a plausible structural basis for the full and partial agonistic activities of 18:1-LPA and 18:0-LPA on LPA1, respectively. Simulations show that 18:1-LPA, in the presence of Gi-protein, retains close interaction between TM3 and TM7 of the receptor, which is not stable in the presence of 18:0-LPA. Chi2 dihedral angles measured in the presence of 18:1-LPA reveal that W271^6.48^ from the C^6.47^W^6.48^xP^6.50^ motif is fully constrained to a rotameric conformation similar to the active LPA1 structure (PDB ID: 7TD1). This further prevents F267^6.44^ from the P^5.50^I^3.40^F^6.44^ motif from leaving its active conformation. However, with 18:0-LPA, TM7 moves away from TM3, possibly due to its limited interaction and the conformation of the activation switch W271^6.48^, which differs considerably in the presence of 18:1-LPA (**Figure 5E**). 18:0-LPA appears to interact more with P273^6.50^ within the C^6.47^W^6.48^xP^6.50^ motif than with W271^6.48^.

Consequently, differences among critical residues in eliciting helical interactions are transferred to the Gi-protein coupling interface. Dynamic network analysis supports the involvement of the N^7.49^P^7.50^xxY^7.53^ motif in the communication path between key binding site residues and the G-protein coupling interface. Specific residues in network paths, including I142^3.46^, R314^7.56^ (18:1-LPA-LPA1), and D315^H8^, S319^H8^ (18:0LPA-LPA1), distinguish communication between the orthosteric site and Gi-protein interface communication in 18:0-LPA- and 18:1-LPA-LPA1 systems. A key consequence at the interface is the differential interaction between K316^H8^ and D350^G.H5.22^ of the G-protein alpha helix, maintained by 18:1-LPA but absent in the 18:0-LPA receptor system. Instead, D350^G.H5.22^ has greater contact with R152^3.56^ in the 18:0-LPA system. The absence of the K316^H8^-D350^G.H5.22^ interaction in the 18:0-LPA-LPA1 system is due to the movement of the TM7 Y311^7.53^ away from the TM3 and its closer interaction with TM1. However, the TM6-TM3 active state conformation is preserved due to strong polar interactions of R146^3.50^ and R152^3.56^ with the Gα helix of Gi-protein. Hence, 18:0-LPA stabilizes a more intermediate conformation of the LPA1 receptor, unlike 18:1-LPA, which stabilizes a fully active LPA1 conformation in the presence of G-protein. Additionally, the presence of Gi-protein is necessary to maintain a fully active conformation of the LPA1, a requirement also noted in β-adrenergic and serotonin (5HT1A) receptors^30, 61^.

WT-metaD simulations provide useful insights into the access paths, critical residue interactions involved, and binding mechanisms of 18:0-LPA and 18:1-LPA to LPA1. The double bond in 18:1-LPA has a moderate influence on its behavior in the aqueous cellular environment, with both ligands having a similar entry process. Both ligands took aqueous paths to access the receptor, with the N-terminal loop playing a crucial role in directing their pre-configuration, orientation, entry, and binding. The presence of free phosphate and hydroxyl moieties makes them more water soluble than other long-chain phospholipids, favoring their access to the orthosteric site via the extracellular aqueous phase after their production from phospholipid precursors^62^. The ligand access through the aqueous route via extracellular vestibules or between transmembrane helices has been demonstrated for endogenous ligands and many xenobiotics in class A GPCRs, including adrenergic receptor and sphingosine-1-phosphate receptor^63,64^. Aqueous origination of ligand entry to LPA1 was earlier proposed due to observations that the spaces between the transmembrane helices of the LPA1 receptor are small for ligand permeation compared to the closely related sphingosine -1-phosphate receptor ^24^. Another study using supervised MD simulations demonstrated that LPA accesses its receptor via the extracellular parts of TM1 and TM7 in a tail-first orientation ^65^. Our simulations show that 18:0- and 18:1-LPA prefer to enter the receptor from the aqueous phase as their phosphate head is anchored by the N-terminal residues R36^Nterm^ and K39^Nterm^ while the highly flexible alkyl chain samples energetically favorable conformations to slide into the receptor via the space between the extracellular loops. These findings align with previous studies, including the mutation of Y34A and K39A, which significantly impaired the ability of LPA1 to couple to Gi-protein, cAMP activation, and intracellular calcium recruitment^66,25, 26, 67^. It should be noted that our simulations do not consider the possible protein-facilitated presentation of the ligand via “chaperones” (e.g., autotaxin)^68^.

We evaluated the membrane partitioning characteristics of 18:0 and 18:1-LPAs using MD simulations. The solvation free-energy profile showed that both ligands have a higher affinity for the membrane lipids than the aqueous phase, with 18:0-LPA showing slightly more favorable partitioning energy than 18:1-LPA **(Figure 11B**). Despite favorable interactions with the bilayer, which serves as a local depot, single-chain lysophospholipids, such as 18:0- and 18:1-LPA, could access their receptor binding site via aqueous routes^63^.

## CONCLUSIONS

This study examined the differential activation, access, and binding mechanisms of 18:0- and 18:1-LPA species to the LPA1 receptor using in vitro and in silico experiments. The results show that 18:1-LPA has higher efficacy for Erk activation through LPA1 than 18:0-LPA in PC-3 human prostate cancer cells. Although the two LPA species possess similar membrane partitioning profiles and access to the orthosteric site of LPA1 through aqueous routes, they have distinct binding modes, residue interactions, and the ability to stabilize the receptor in active state conformations. Notably, the presence of a double bond in 18:1-LPA exerts conformational restraints within the binding site, promoting distinct interactions with the activation switch residues, W271^6.48^ and F267^6.44^, stabilizing the receptor conformations in an active state in the presence of Gi protein. In contrast, 18:0-LPA has less contact with the activation residues, resulting in less stabilization of the receptor and distinct Gi protein interactions, possibly explaining its partial agonism profile. Despite their energetically favorable membrane partitioning profiles, both molecules take aqueous paths to access the orthosteric site, revealing the functional role of N-terminal residues in promoting access, pre-organization, and binding of LPA species through specific residue-ligand interactions. In summary, this study provides valuable insights into the structural basis and ligand-receptor dynamics underlying the access, binding, and differential activation mechanisms of two LPA species on LPA1.

## MATERIALS AND METHODS

### In Vitro Studies

#### Materials

LPA species were obtained from the following sources: (18:1; oleoyl) from Echelon Biosciences (Salt Lake City, UT), and LPA (18:0; stearoyl) from Avanti Polar Lipids (Birmingham, AL). LPAs were prepared in a stock solution with 4 mg/ml fatty-acid bovine serum albumin (BSA), stored at -20°C, and delivered to cells as a 1000X stock solution. Vehicle controls for LPA had a final concentration of 4 µg/ml BSA. Anti-phospho-Erk (#9106) was purchased from Cell Signaling Technologies (Danvers, MA) and used at 1:1000 dilution. Anti-GAPDH (sc-25778), obtained from Sant Cruz Biotechnologies (Dallas, TX), was used at 1:2000 dilution. Anti-rabbit IgG secondary antibody (#Lot 31) and Anti-mouse IgG, HRP-linked Antibody (#7076) were purchased from Cell Signaling Technologies (Danvers, MA) and used at 1:2000 dilution.

#### Cell culture

PC-3 cells were obtained from the American Type Culture Collection (Manassas, VA). The cells were grown in RPMI 1640 medium (Cytiva, Marlborough, MA) with 10% FBS (VWR, Visalia, CA), on standard tissue culture plastic in an incubator maintained at 37°C with 5% CO2.

#### Immunoblotting

Cells were rinsed twice with ice-cold phosphate-buffered saline (PBS), harvested by scraping into 1 ml ice-cold PBS, collected by centrifugation at 10,000xg for 10 min at 4°C, and resuspended in ice-cold lysis buffer (20 mM HEPES [pH=7.4], 1% Triton X-100, 50 mM NaCl, 2 mM EGTA, 5 mM β-glycerophosphate, 30 mM sodium pyrophosphate, 100 mM sodium orthovanadate, 1 mM phenylmethylsulfonyl fluoride, 10 µg/ml aprotinin, 10 µg/ml leupeptin). Insoluble debris was discarded after centrifugation. Whole-cell extracts containing equal amounts of protein (30µg) were separated by SDS-PAGE on 12.5% Laemmli gels, transferred to PVDF membranes presoaked in 100% methanol, and incubated with primary (overnight at 4 °C) and then secondary (one to two hours at room temperature) antibodies. Blots were developed using enhanced chemiluminescence reagents (GE Healthcare) and imaged using a Gel Doc system (BioRad Laboratories, Hercules, CA). Protein expression was quantified by densitometry using Image J software. Results were normalized to the GAPDH loading control and then to the value obtained for untreated control cells to evaluate the fold increase.

### In Silico Studies

#### Protein and ligand structure preparation

In this study, we used the active structure of LPA1-Gi-protein complex bound to 18:1-LPA (PDB ID: 7TD1). The structure preparation step was done using the MOE software. The *QuickPrep* module in MOE was utilized to generate rotamers, assign appropriate protonation states to residues, add the missing hydrogen atoms, and minimize the structure to remove any bad contacts. All the titratable residues were assigned their dominant protonation states at pH 7.4 using the GBSA solvation model. The 3D atomic coordinates of 1-stearoyl-2-hydroxy-*sn*-glycero-3-phosphate (18:0-LPA) were obtained from PubChem website ^20^. The geometries of the ligand structures were further optimized using MOE.

#### Molecular Docking

The molecular docking of 18:0-LPA at the orthosteric binding site of the receptor was done using the docking module in MOE^69^. The binding pocket was defined by select residues, including R124^3.28^, D129^3.33^, W210^5.43^, G274^6.51^, and K294^7.36^ within the orthosteric site of the receptor. We used Alpha PMI and Triangle matcher placement methods to generate multiple poses of the ligands within the receptor. The protein-bound ligand structures were further refined using the induced-fit method, in which the receptor side chains were allowed to be flexible while the backbone was fixed. The docked poses were then ranked using the London dG scoring function. The most appropriate poses were selected based on the docking score, observed interactions, and orientation of the functional groups of the ligand within the orthosteric site.

#### Unbiased MD Simulations

The four receptor-ligand complexes (18:1-LPA-LPA1-Gi protein, 18:1-LPA-LPA1, 18:0-LPA-LPA1-Gi protein, 18:0-LPA-LPA1) were then oriented in the membrane plane using the OPM server^70^. The membrane builder module of the CHARMM-GUI web server was utilized to generate a heterogenous membrane consisting of 16:0/18:1 phosphatidylcholine (POPC) (80-90) and cholesterol (8-11) ^71^. Ligand parameterization was carried out using CHARMM-GUI ligand reader and modeler, and charges were assigned using CGenFF ^72^. The system was solvated with the TIP3P water model padding up to about 23 Å on both sides. The ionic concentration of the final system was at 0.15 M using sufficient NaCl. The final ligand-receptor-membrane complex, which contained 205 lipid molecules, was then subjected to 5000 steps of minimization and thereafter was equilibrated using the CHARMM-GUI recommended six-step equilibration protocol^71, 73^. The equilibration steps involve applying various restraints to the system components: 1) harmonic restraints to ions and heavy atoms of the protein, water, ion, and lipid molecules, 2) repulsive planar restraints on water to prevent them from entering into the bilayer core region, and 3) planar restraints to hold the lipid headgroups in position along the bilayer normal. During the initial equilibration step, the harmonic restraints were applied to all the system components at their maximum levels [10, 5, 2.5, 2.5, and 10 kcal/(mol*Å^2^) on protein backbone atoms, protein sidechain, water, lipid and ions, respectively]. In the subsequent steps, the restraints were removed from ions, and the force constants were gradually reduced for other components. Complete details are provided in Table S5^71, 73^. All MD simulations were run using GROMACS 5.1.4 suite ^74^. During the production run, van der Waals and short-range electrostatic interactions were estimated with a cut-off distance of 12 Å. The long-range electrostatic interactions were computed using the particle-mesh Ewald summation method. A 2-fs integration time step was used for all MD simulations. The system was simulated under conditions of constant pressure and temperature (NPT conditions) at 1 atm pressure and temperature of 310 K, respectively. The temperature and pressure were controlled using the Nose-Hoover thermostat and the Parinello-Rahman barostat with a coupling constant of 5.0 ps and compressibility of 4.5x10^-5^ bar^-1^. All simulations were carried out for a duration of 500 ns with three replicates per simulation system. Frames from the simulation were saved every 10 picoseconds.

#### Membrane partitioning simulations

The membrane partitioning characteristics of 18:0-LPA and 18:1-LPA were investigated using enhanced sampling methods, including steered molecular dynamics (SMD) and umbrella sampling (US), as previously described^75^. The CHARMM-GUI ligand reader and modeler were used for ligand parameterization, and charges were added using the CGenFF^76^. The CHARMM-GUI membrane builder module was utilized to assemble the ligands and the lipid bilayer ^77–79^. The bilayer composition in both 18:1-LPA and 18:0-LPA simulations included 70 POPC molecules and seven cholesterol molecules in each of the upper and lower leaflets. The entire system was solvated using the TIP3P water solvation model, while an ionic concentration of 0.15 M NaCl was maintained ^80^. The CHARMM36 force field parameters were utilized to model interactions among the system components ^79, 81^. Subsequently, the ligand/bilayer system was subjected to equilibration following the CHARMM-GUI protocol, which involves six steps in total. The first two steps of the equilibration utilize the NVT ensemble, while the last four steps make use of the NPT ensemble. Certain restraints were introduced at the start of the equilibration. A harmonic force constant of 5 kcal/mol was applied on the heavy atoms, planar restraints, which restrain the lipid head groups of the bilayer components along with the z-axis, and dihedral restraints to limit the chirality of double bonds and lipid headgroups. This lasted for 250 ps. The following two steps included a scaling factor on the restraints and subsequent stepwise release of the restraints, which also lasted for 250 ps. The last three steps of the equilibration protocol lasted 500 ps and included a scaling factor on the restraints. However, the restraining force was removed entirely during the last step. Thereafter, an additional equilibration run was executed for 50 ns before the beginning of the SMD process. At the start of the SMD simulation, the ligand was kept within the aqueous phase at 30 Å from the center of the membrane bilayer. SMD was used to pull the ligand from its position in the aqueous bulk through the membrane bilayer along the bilayer normal (z-axis). The pulling occurred at 1 Å per ns rate, integrated by a one fs time step with a harmonic force restraint of 5 kcal/mol/Å on the LPA molecule. The ligand/bilayer system coordinates at each 1 Å of the permeability path (totaling 30 windows) were obtained from the SMD simulation and used as starting structures for the subsequent umbrella sampling simulations. For the umbrella sampling, the system representing each window was first equilibrated for 10 ns, and a subsequent production simulation was run for an additional 50 ns. This resulted in a cumulative simulation time of ∼1.5 µs for each of the ligands studied. The z-component of the distance between the center-of-mass of the lipid atoms and the heavy atoms of the ligand was restrained by a harmonic force of 1.5 kcal/mol/Å during the umbrella sampling simulations. This was implemented through the Colvars module. ^82^ The obtained probability distributions were reweighted using the weighted histogram analysis method (WHAM) ^83^ to derive an unbiased Potential Mean Force (PMF). All SMD and US simulations were run using the CUDA version of NAMD 2.12.

#### Association Simulations

Metadynamics has evolved as an excellent tool for studying rare and important events within realistic timescales^84^. In metadynamics, a history-dependent external potential acts on a few degrees of freedom, known as collective variables. This imposes an energetic barrier, consequently limiting the system from revisiting already sampled configurations. Well-tempered metadynamics (WT-MetaD) is technique that applies restraint on the height of the bias potential in relation to the amount of bias already deposited. Hence, it is well suited to sample the free-energy surface (FES) for the ligand access to its receptor ^85^. Using WT-MetaD, we investigated the potential access route to the orthosteric site for both 18:0-LPA and 18:1-LPA starting from the aqueous phase and energetically favorable locations within the membrane bilayer. The system was equilibrated using the CHARMM-GUI-based 6-step protocol.

The WT-MetaD simulations were carried out using GROMACS 2019 ^74, 86^ patched with PLUMED v2.3^87, 88^. Two collective variables (CV) were chosen for the association process. The first CV was the distance between the center-of-mass (COM) of the binding site residues, G110^ECL1^, R124^3.28^, W210^5.43^, E293^7.35^, and COM of the ligand. The second CV was the internal angle of the ligand, represented by three points, namely the phosphate head group, the upper part of the alkyl chain (C9-C17), and the tail part of the alkyl chain (C1-C8) **(Figure 2A)**. The simulation parameters included a temperature of 310K, an upper wall boundary of 3.0 nm, a height of 1.5, a sigma value at 0.05, a bias factor of 15, and a force of 200 kJ/mol. The resulting free-energy surface was generated using the MEPSA tool ^89^. All simulations were carried out in triplicates.

#### Network Analysis

Network analyses of the trajectories from the unbiased simulations were performed to investigate the potential signal transduction paths between the activation site residues (R124, W271, and K294) and the residues that interface with Gi-protein (R152 and K316). In a typical receptor network model, the nodes (represented by amino acid residues) are connected by edges. Two nodes are in contact if their heavy atoms are within 4.5 Å for over 75% of the simulation. The length of a path between two distant nodes is the sum of the edge weights between the consecutive nodes along the path. The shortest path (optimal path) between the distant nodes is assumed to be their most prevalent communication path. The dynamic network analyses between the chosen nodes (residues) were calculated using the *networkView* plugin in VMD software ^90–92^. The contact maps between amino acid residues were obtained using the *Carma* software ^93^. The communities of closely associated residues were generated using the *gncommunities* module, while the *subopt* program was used to identify suboptimal paths between the residues of interest.

#### Binding Free-Energy Calculations

The Molecular Mechanics Poisson-Boltzmann Surface Area (MMPBSA) technique was utilized to calculate the binding free energy of the two ligands. We also calculated the relative energy contribution attributable to each residue. The MMPBSA technique utilizes molecular mechanics force field parameters to derive the average molecular mechanical energy and continuum solvation models ^94, 95^. The MMPBSA method was applied using the most recent version of the gmx_MMPBSA tool^96^. The average molecular mechanical energy (Δ*E*_MM_) is the summation of both bonded (Δ*E*_bond_) (bond, angle, dihedral, and other internal energies), electrostatics (Δ*E*_ele_), and van der Waals (Δ*E*_vdW_) interaction energies. The calculations were performed using the molecular mechanics (MM) force-field parameters Δ*E*_MM_ can be summarized as the following: Δ*E*_MM_ = Δ*E*_ele_ + Δ*E*_vdW_ + Δ*E*_bonded_

The solvation model takes into consideration polar (Δ*G*pb) and nonpolar (Δ*G*np) solvation free energies and the solvent-accessible surface area (SASA). The energy contribution of each protein residue was then derived using the following equation ^97^: where Δ*R^BE^* is the binding energy of residue *x*, *n* is the total amount of atoms in the residue, and *A^free^* are the energy of *i*th atom from *x* residue in bound and unbound states, respectively.

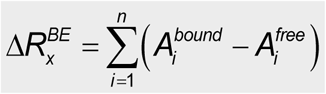

## Data Availability

The data used to generate results, including starting structures, simulation parameters, and trajectory files, can be accessed at:https://tinyurl.com/3hcfkps7 or https://doi.org/10.5281/zenodo.14796886

## Supporting Information

Supporting Information is available free of charge at:

- Physicochemical properties of LPA species; Dihedral angle profile of key aromatic residues in experimental LPA1 structures; Membrane partitioning profile of LPA species; Binding poses of the LPA species in LPA1; Ligand stability and internal angle; Contact fingerprint of the ligands’ alkyl chains; LPA1-Gi-protein contact analyses; Comparative analyses of important activation signatures in different LPA1-ligand systems; Sequence alignment of the six LPA receptors. (PDF)
- **Movie M1** showcases the binding pose and molecular interactions of 18:1 LPA at the orthosteric site of the LPA1 receptor observed in all-atom classical MD simulations.
- **Movie M2** depicts the binding modes and important residue interactions of 18:0 LPA with the LPA1 receptor observed in all-atom MD simulations.
- **Movie M3** depicts the interaction between 18:1-LPA-bound LPA1 and the α5 helix of the Gα subunit of Gi-protein. The conserved R146^3.50^ forms a stable hydrogen bond interaction with C351 (from Gαi protein)
- **Movie M4** depicts the receptor-G-protein interface interaction between 18:0-LPA-bound LPA1 and the α5 helix of the Gα subunit of Gi-protein.
- **Movie M5** shows the access and binding processes of 18:1 LPA to LPA1 from the extracellular aqueous environment. This association simulation was performed using well-tempered metadynamics.
- **Movie M6** illustrates the access and binding processes of 18:0 LPA to the LPA1 receptor from an extracellular aqueous environment.

## Notes

The authors declare no competing financial interests.

---

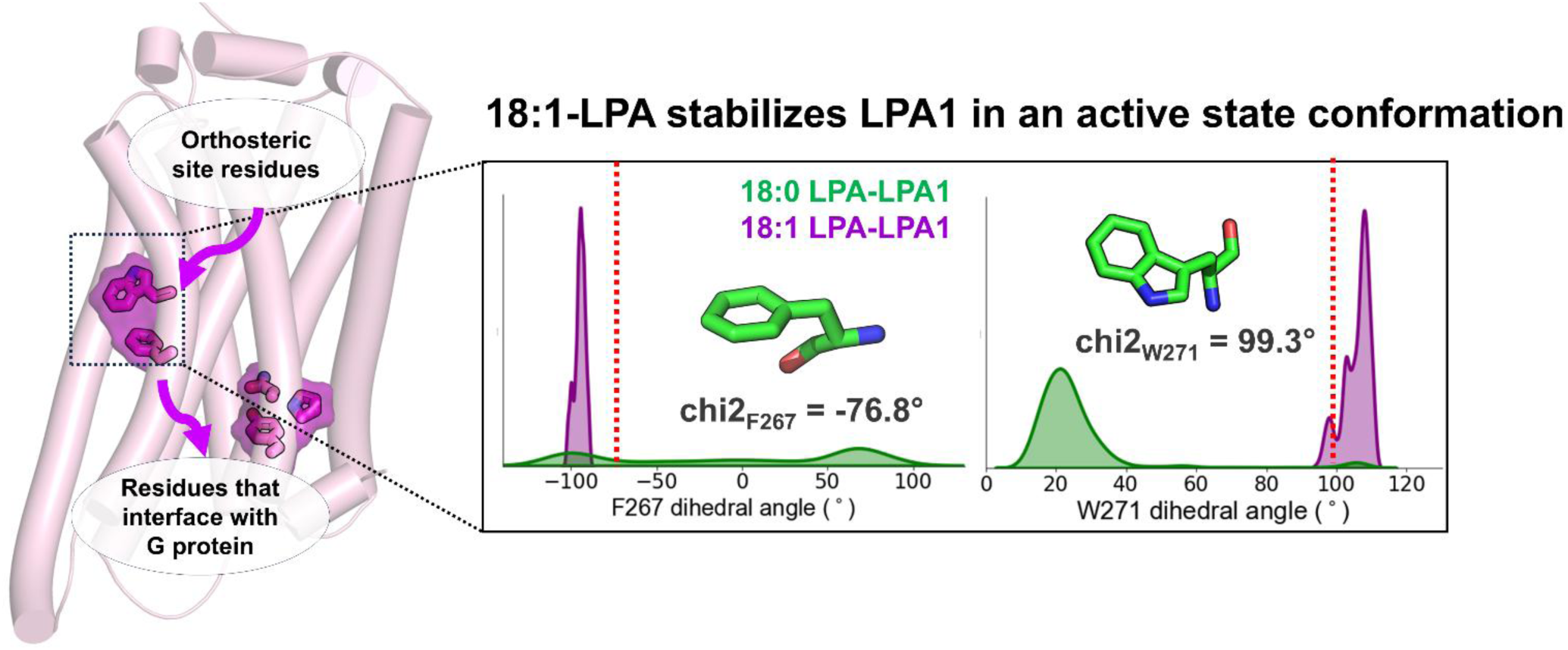

